# Spatial proteomics reveals CD8+ T cell signatures and cellular niches associated with active HIV-1 replication in lymph nodes

**DOI:** 10.64898/2026.05.11.724393

**Authors:** Candace C. Liu, Marta Calvet-Mirabent, Angie Spence, Mako Goldston, Magdalena Adrados de Llano, Sean C. Bendall, Michael Angelo, Enrique Martín-Gayo

## Abstract

Despite its effectiveness in suppressing active HIV-1 replication, antiretroviral therapy (ART) does not eliminate the persistent long-lived pool of HIV-1-infected reservoir cells, preventing the eradication of the infection. Lymphoid tissues are key anatomical sites where these reservoirs persist even in the presence of ART, but the mechanisms that are associated with viral persistence in lymphoid tissues and how tissue networks are reshaped in the setting of viral replication remain incompletely understood. Advances in tissue imaging offer a unique opportunity to characterize immune correlates of viral persistence. Here, we used a spatial proteomic method to map immune microenvironments in HIV-1-infected lymph nodes (LNs) at different stages of infection, including with or without ART. LNs from people with HIV-1 (PWH) were characterized by lower CD4+ T cell counts and higher CD8+ T cell counts in both the whole tissue and within follicles compared to people without HIV (PWOH). CD8+ T cells were more abundant in LN samples with active viral replication, defined by detection of the viral protein p24. Further characterization of p24+ LNs showed that CD8+ T cells located inside of B cell follicles exhibited higher levels of markers associated with immune activation and exhaustion, in addition to the inflammasome protein caspase-1. Using a spatial niche detection method, we found that LNs from PWH with varying levels of viral replication were differentially enriched for CD8+ T cells near antigen-presenting cells, myeloid cells, and fibroblasts. Notably, we found that p24+ cells were less enriched near CD8+ T cells but closer to follicular dendritic cells. Finally, comparing LNs from viremic and aviremic donors, where viremia was defined by detectable plasma viral load, we found low levels of activation markers in CD11c+ cells in aviremic donors, including NLRP3 inflammasome activation. Thus, using spatial proteomics to map the immune landscape in LNs, we identified novel markers characterizing immune cell subsets and tissue microenvironments that were differently enriched in PWH with varying levels of viremia, implying that HIV-1 infection confers long-term changes on the immune landscape in LN tissue. Collectively, these data provide new insights into the complex cell networks associated with viral replication at a key tissue reservoir site, which could be relevant for future HIV-1 cure strategies.

## Introduction

HIV-1 infection remains one of the most significant global health challenges, with over 40 million people living with the virus worldwide^1^. While the introduction of antiretroviral therapy (ART) has transformed HIV-1 infection from a fatal disease to a manageable chronic condition, achieving a functional cure remains elusive due to the persistence of latent viral reservoirs in different tissues^2–7^. These reservoirs are composed of heterogeneous infected cells, which can reactivate HIV-1 replication and produce infectious viral particles when ART is discontinued, necessitating lifelong treatment for people with HIV-1 (PWH)^2,8,9^.

While most previous studies on HIV-1 reservoirs have focused on peripheral blood, lymph nodes (LNs) serve as critical anatomical sites for HIV-1 pathogenesis and persistence^4,10–15^. The complex architecture of LNs, with distinct anatomical compartments including B cell follicles, germinal centers, and T cell zones, creates specialized niches that may influence the dynamics between reservoir establishment, viral replication, and inflammatory responses. Understanding the spatial organization of immune responses within these tissues is essential for elucidating the altered landscapes that could be limiting the efficacy of current immunotherapy candidates and for improving cure strategies.

While CD8+ T cells play a central role in controlling HIV-1 infection through their ability to recognize and eliminate virally infected cells^16–18^, their effectiveness in LNs is limited by several factors, including the potential development of T cell exhaustion, which is characterized by the progressive loss of effector functions and increased expression of inhibitory receptors, commonly observed in chronic viral infections. Our previous work indicated that memory CD8+ T cells from PWH with different checkpoint receptor patterns have different functional profiles, and PD1, TIGIT, and TIM3 are associated with the ability to eliminate HIV-1-infected CD4+ T cells^19^. Viral replication has also been associated with inflammatory pathways such as inflammasome activation that may affect the immune landscape in LNs^20–22^.

In addition, the “follicular sanctuary” hypothesis proposes that infected cells within B cell follicles are protected from CD8+ T cell-mediated killing due to the exclusion of CD8+ T cells from these regions, allowing viral reservoirs to persist in follicular helper T (Tfh) cells and other populations^23– 25^. Studies in primate and human tissues suggest that immune control of HIV-1 infection may correlate with the capacity of cytotoxic CXCR5+ CD8+ T cells to infiltrate into B cell follicles^26–32^. While it has previously been described that a higher proportion of cytotoxic follicular CD8+ T cells is associated with viral control, the functional characteristics, including the exhaustion and inflammatory state of these infiltrating cells, are not fully understood. Moreover, our understanding of the spatial relationships between these effector cells and HIV-1-infected cells in lymphoid tissue remains incomplete.

Here, we present the first comprehensive spatial proteomic analysis of LNs from PWH using multiplexed imaging by time-of-flight (MIBI-TOF)^33,34^. By simultaneously profiling 40 proteins across HIV+ and HIV-donors, we characterized the fundamental alterations in LN architecture associated with HIV-1 infection and their relationship to viral replication status. We found substantial CD8+ T cell infiltration in HIV+ LNs, including into B cell follicles, with these cells exhibiting distinct activation states in tissues showing viral activation, including higher expression of the inflammasome protein caspase-1 and differential expression patterns associated with checkpoint, cytotoxicity, and effector function. We found that the spatial proximity of CD8+ T cells to antigen-presenting cells (APCs) and stromal populations varied based on viral replication status. Additionally, we observed altered myeloid cell compositions and lower inflammasome activation in tissues from aviremic PWH. These findings provide a rationale for how distinct spatial niches within LNs, characterized by elevated checkpoint receptor expression, inflammasome activation, and CD8+ T cell association with myeloid and stromal cells, may contribute to HIV-1 persistence.

## Results

### Mapping the global immune landscape in lymph nodes

To investigate the changes in cellular organization in LNs due to HIV-1 infection, LN tissue from five PWH were obtained from the Pathology Anatomy Unit at Hospital Universitario La Princesa (Madrid, Spain), in addition to LNs from five people without HIV-1 (PWOH) included as negative controls (Figure 1a). Among the PWH, two donors were individuals on ART with fully suppressed viremia, while three were individuals with detectable plasma viral loads (Figure S1a, Supplementary Table 1). Of the latter group, two tissue samples exhibited detectable staining of the HIV-1 protein p24, which serves as a marker for active viral replication. The two samples with detectable p24 tissue staining correspondingly had the highest plasma viral loads (Supplementary Table 1). Out of these two samples, one donor was sampled at the time of diagnosis, and the other donor had discontinued ART about four months prior to biopsy.

**Figure 1:**
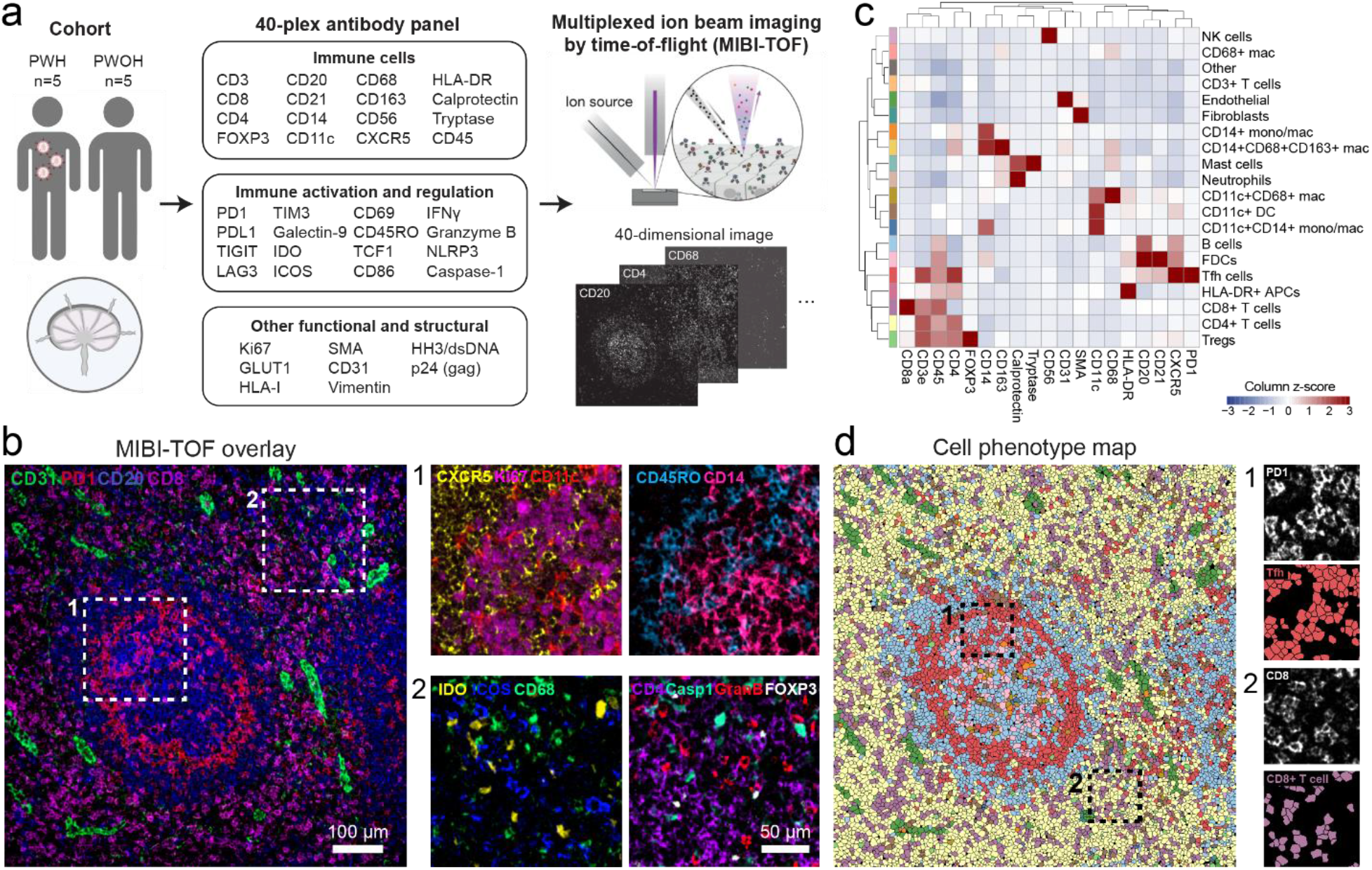
MIBI-TOF imaging reveals the cellular landscape of lymph nodes from PWH and PWOH. (a) Study overview. PWH: people with HIV, PWOH: people without HIV. (b) Representative MIBI-TOF overlays of a single field-of-view (FOV) (800 µm x 800 µm). (c) Heatmap of cell phenotypes identified across the cohort, with identified phenotypes in the rows and markers in the columns. Values were z-score scaled for each marker. (d) Corresponding cell phenotype map of FOV shown in b, where single cells are colored according to the color bar in c.

To deeply characterize these LN samples, we applied MIBI-TOF to simultaneously image 40 proteins in the tissue, capturing multiple anatomical regions within each LN, including T cell zones, B cell follicles, and germinal centers (Figure S1b). The antibody panel included markers for immune cell and stromal lineages, immune cell activation, exhaustion, and inflammation, as well as the viral antigen p24 (Figure 1a-b, Figure S2). Following image acquisition, we employed a custom computational pipeline to profile over 1.7 million cells across the cohort (Figure 1c-d, Figure S3, Figure S4a-b). We assigned the cells to 20 different immune and stromal lineages, characterized the expression of various functional and inflammatory markers, and mapped the spatial relationships between these cell types within their native tissue context (Figure S3, Methods). Altogether, we created a comprehensive cell atlas of LNs from PWH and PWOH to serve as a foundation for investigating virus-associated changes in tissue structure.

### HIV-1 infection shapes CD8+ T cell functional profiles in LNs

To systematically identify differences between LNs from PWOH (HIV-) and PWH (HIV+), we quantified over 1000 features for each image, encompassing cell density and functional marker expression, as well as spatial characteristics including cellular diversity, cellular microenvironments, and cell-cell enrichment (Figure 2a, left). Using a mixed-effect model to identify the features most enriched in HIV- or HIV+ LNs, we found that despite varying treatment status and viremia levels, HIV+ LNs consistently exhibited fewer CD4+ T cells and more CD8+ T cells compared to HIV-LNs (Figure 2a, right). The CD4+ to CD8+ T cell ratio perfectly segregated donor groups (Figure 2b). Notably, LNs from all PWH displayed a lower CD4+ to CD8+ T cell ratio than PWOH, including samples from PWH donors with undetectable plasma viremia and absence of tissue p24 staining, suggesting that the presence of virus induces long-lasting changes to LN architecture that persist even after viral suppression (Figure 2b). Interestingly, we found that there was little correlation between the CD4+ or CD8+ T cell counts in the LN tissues and corresponding blood counts measured at the time of tissue sampling (Figure S4c).

**Figure 2:**
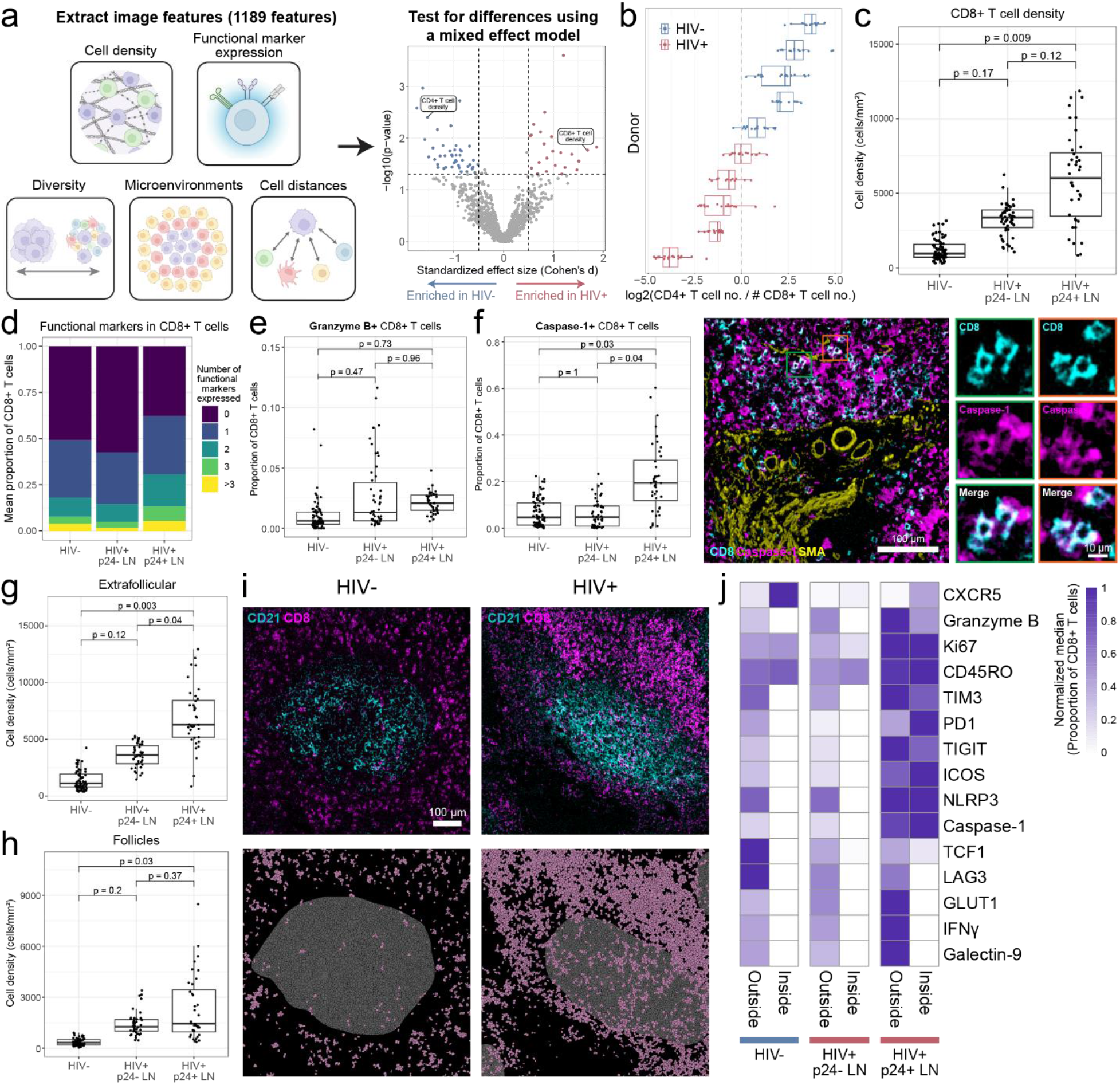
Viral replication status shapes CD4+ and CD8+ T cell phenotypes in LNs from PWH. (a) Overview of feature extraction pipeline. A mixed-effect model was used to test for significance between HIV+ and HIV-LNs for all features. (b) The log2 ratio of CD4+ T cells and CD8+ T cells for all donors. Each dot is one FOV. (c) Comparison of CD8+ T cell density in HIV-donors, HIV+ donors where p24 was undetectable, and HIV+ donors where p24 was detected in any FOV of the LN from that donor. Each dot is one FOV. (d) Breakdown of the proportion of CD8+ T cells that express functional markers. Functional markers include caspase-1, CD45RO, CD69, galectin-9, GLUT1, granzyme B, ICOS, IFNg, Ki67, LAG3, NLRP3, PD1, PDL1, TCF1, TIGIT, TIM3. (e) Comparison of proportion of CD8+ T cells that are positive for granzyme B and (f) caspase-1. Representative MIBI-TOF images shown on the right. (g) Comparison of the density of CD8+ T cells in the extrafollicular or (h) follicular areas. (i) Two representative FOVs. Top: MIBI-TOF images, bottom: cell phenotype maps (gray: follicle area, pink: CD8+ T cells). (j) Normalized median proportion of CD8+ T cells expressing functional markers, inside or outside the follicle area.

Further stratifying the donors by p24 detection in tissues (Figure S5a), we found a significant increase (p = 0.009) in CD8+ T cell density specifically in LN from PWH where p24 was detected in any imaged area of the tissue (p24+ LNs, Figure 2c). LNs from PWH without detection of p24 (p24-LNs) were characterized by an intermediate phenotype between p24+ LNs and HIV-LNs, exhibiting a moderate expansion of CD8+ T cells. In parallel, p24+ LNs showed significant CD4+ T cell depletion (p = 0.02, Figure S5b). Interestingly, the most significant feature of HIV-LNs from the mixed-effect model was a higher density of TCF1+ CD4+ T cells (Figure S5c), a cell subset that is associated with stemness and memory cell differentiation^35,36^. In HIV+ LNs, the most significant feature was a higher spatial enrichment score between CD11c+ CD14+ myeloid cells and CD4+ T cells, indicating that the distribution of these myeloid cells and CD4+ T cells was influenced by viral replication (Figure S5d). These data demonstrate that active viral replication, as evidenced by p24 expression, is associated with substantial changes in tissue structure in both CD4+ and CD8+ T cell populations.

Given the elevated CD8+ T cell counts in HIV+ LNs relative to HIV-control tissues, we hypothesized that CD8+ T cells in p24+ versus p24-LNs would exhibit different inflammatory and exhaustion signatures. Surprisingly, we found that a large proportion of the CD8+ T cells lacked any of the functional markers associated with immune activation, exhaustion, and inflammation included in our panel (CD45RO, CD69, galectin-9, GLUT1, granzyme B, ICOS, IFNγ, Ki67, LAG3, NLRP3, PD1, PDL1, TCF1, TIGIT, TIM3, caspase-1) in p24-LNs (Figure 2d). In fact, most CD8+ T cells expressed none or only one of these markers, with few cells expressing multiple markers simultaneously. Expectedly, p24+ LNs contained the highest proportion of CD8+ T cells expressing one or more of these functional markers, indicating higher levels of CD8+ T cell activation in response to viral protein expression (Figure 2d).

Further analysis of activated CD8+ T cells revealed distinct functional patterns across donor groups. While not statistically significant, the proportion of CD8+ T cells that expressed the cytotoxic molecule granzyme B trended higher in HIV+ LNs, including both p24- and p24+ tissues, suggesting an increase of cytotoxicity regardless of active viral replication (Figure 2e, Figure S5e). Notably, we observed a significant increase (p = 0.03) in caspase-1+ CD8+ T cells specifically in p24+ LNs (Figure 2f, Figure S5f). While caspase-1 expression is typically associated with inflammasome activation and pyroptotic cell death in myeloid cells, its expression in CD8+ T cells has been less well characterized. Furthermore, we found that the proportion of granzyme B+ caspase-1+ double-positive CD8+ T cells trended higher in p24+ LNs, while granzyme B+ caspase-1-CD8+ T cells trended higher in all HIV+ LNs (Figure S5g-j). The expression of other markers associated with T cell activation, such as the proliferation marker Ki67, memory marker CD45RO, effector molecule IFNγ, and checkpoint molecules TIGIT, TIM3, and ICOS, also trended higher in CD8+ T cells from p24+ LNs (Figure S5k-l). Interestingly, expression of galectin-9, the ligand for TIM3, was also expressed higher in CD8+ T cells from p24+ LN (Figure S5k-l).

To gain deeper insight into the functional implications of the phenotypic differences in CD8+ T cells, we analyzed functional marker co-expression patterns. Assessment of the most frequently co-expressed functional marker pairs in CD8+ T cells revealed distinct differences between donor groups (Figure S5m). For example, caspase-1+ CD45RO+ co-expression was dominant in HIV+ p24+ LNs and HIV-LNs, while Ki67+ CD45RO+ co-expression was most common in p24-LNs. These data suggest different inflammatory versus proliferative phenotypes in memory CD8+ T cells in LN with active viral replication. While CD8+ T cells in HIV+ p24-LNs showed co-expression patterns dominated by CD45RO, CD8+ T cells in p24+ LNs exhibited increased checkpoint molecule co-expression. Focusing on the checkpoint markers, while we found that the majority of CD8+ T cells did not express any of these markers, the most common co-expression pattern in CD8+ T cells was TIGIT and PD1, followed by cells co-expressing TIM3 and galectin-9, and TIGIT and TIM3 (Figure S5n-o). Specifically looking at granzyme B, which was increased in all HIV+ LNs, and caspase-1, which was increased in p24+ LNs, we found that both markers were highly co-expressed with CD45O (Figure S5p-q). Analysis of granzyme B+ or caspase-1+ CD8+ T cells revealed minimal co-expression with checkpoint markers, suggesting that upregulation of these markers is not concurrent with immune exhaustion in these cells (Figure S5r). Together, these findings demonstrate that CD8+ T cells in HIV+ LNs exhibit a spectrum of activation states, showing increased inflammatory and exhaustion profiles in p24+ tissues.

### CD8+ T cell infiltration into B cell follicles

To determine whether the observed CD8+ T cell phenotypes in LNs were associated with exclusion from B cell follicles, we analyzed the distribution and phenotype of these cells in extrafollicular versus follicular areas. We found an increase in CD8+ T cells in the extrafollicular areas of HIV+ LN, which was significantly higher (p = 0.003) in p24+ LNs (Figure 2g). Interestingly, CD8+ T cell densities were elevated within follicular regions across all HIV+ LNs, with significantly higher levels (p = 0.03) in p24+ LNs (Figure 2h-i). These data indicated that CD8+ T cells can access anatomical compartments previously described to harbor viral reservoirs, regardless of active viral replication, but tend to accumulate outside follicles in the absence of suppression.

To further characterize patterns associated with active viral replication, we then analyzed the phenotypic differences between CD8+ T cells located in extrafollicular or follicular areas in the different donor groups. We confirmed that granzyme B expression was generally higher in HIV+ LNs, but particularly in cells located in the extrafollicular areas (Figure 2j). In contrast, Ki67 and CD45RO were higher in p24+ LNs, with minimal differences between follicular and extrafollicular areas. Interestingly, the expression of various checkpoint and activation markers, including TIM3, PD1, TIGIT, and ICOS, as well as inflammasome markers NLRP3 and caspase-1, were elevated in the follicles specifically in p24+ LNs, and low in both p24- and HIV-LNs. This marker profile indicates that follicle-infiltrating CD8+ T cells represent activated, proliferating populations that may be responding to local antigen presentation or microenvironmental cues. In contrast, CD8+ T cells that expressed TCF1, LAG3, GLUT1, IFNγ, and galectin-9 were largely excluded from the follicles (Figure 2j). Collectively, this data suggests that HIV-1 infection and active viral replication, as well as histological localization, affect the activation and exhaustion states of CD8+ T cells.

### CD8+ T cells exhibit distinct spatial niches in HIV+ LNs with active viral replication

Cellular function is strongly influenced by the characteristics of the local tissue microenvironment. To identify differential spatial relationships between HIV- and HIV+ LNs, we used QUICHE, a graph-based spatial niche detection method, where niches refer to groups of cell types in close proximity in the tissue^37^. Using this unbiased approach without prior specification of CD8+ T cells, QUICHE identified spatial relationships involving CD8+ T cells as key distinguishing features of HIV+ LNs (Figure 3a). The niche defined by CD8+ T cells nearby other CD8+ T cells showed the largest mean fold change difference between the HIV+ and HIV-groups, with a higher value in the HIV+ group. Furthermore, we found that HIV+ LNs were enriched for CD8+ T cells in close proximity to APCs (HLA-DR+, CD11c+, CD14+), fibroblasts, and endothelial cells. We confirmed this finding using cell enrichment analysis (Figure S6a-b). In contrast, HIV-LNs showed enrichment for CD4+ T cell and Tfh cell niches. Interestingly, we found that the significantly enriched niches in the HIV+ group were mostly capturing interactions outside of the follicles, with niches containing CD11c+ CD14+ myeloid cells present to a limited extent in the follicles (Figure S6c). The most abundant niches within follicles consisted of expected cell types, predominantly B cells, follicular dendritic cells (FDCs), and Tfh cells (Figure S6d).

**Figure 3:**
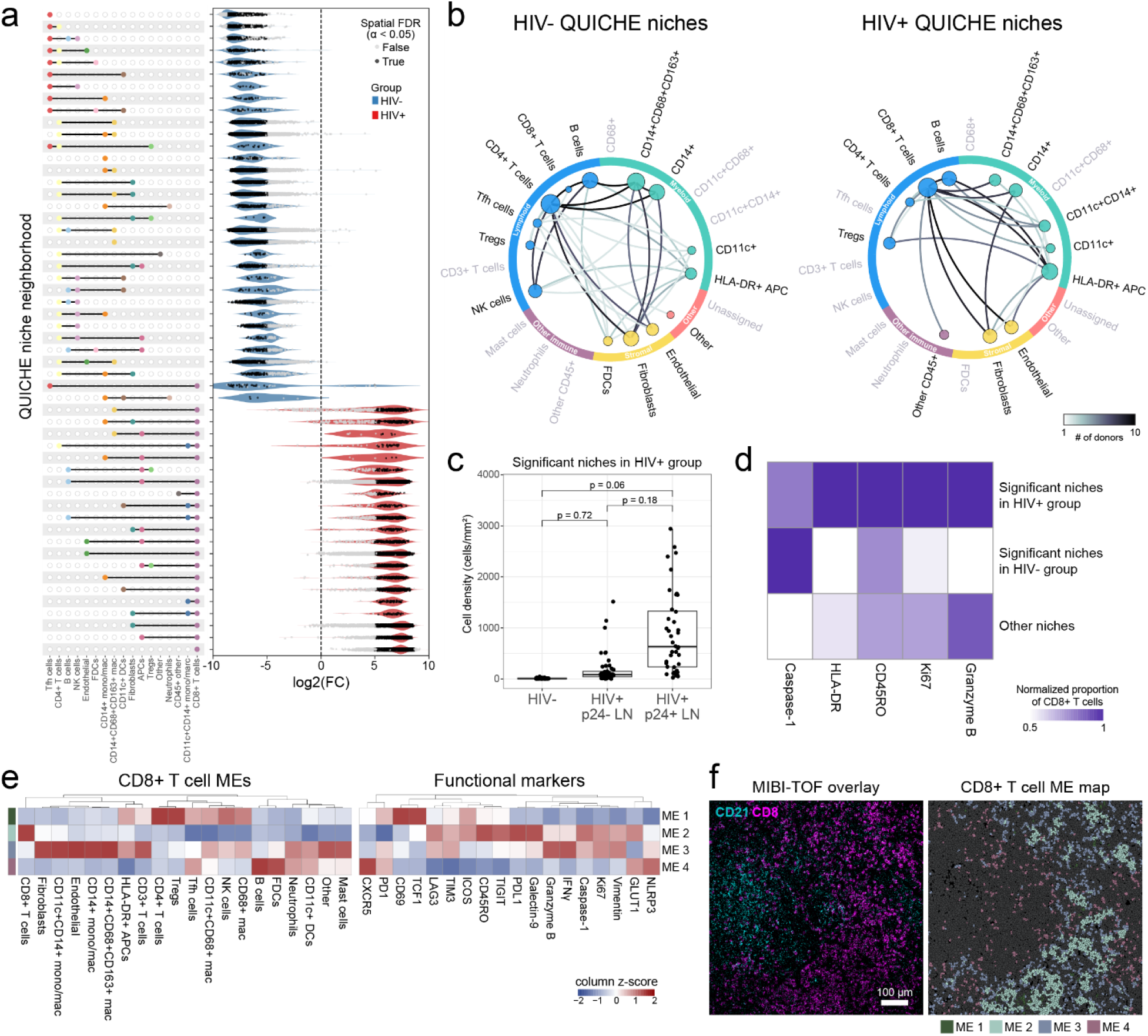
Spatial analysis identifies distinct CD8+ T cell niches associated with HIV-1 infection and active viral replication. (a) Violin plots show the top differentially enriched niche neighborhoods within HIV-(blue) or HIV+ (red) LN samples as identified using QUICHE. Niches were filtered for median logFC > 1 or < -1 and median Spatial FDR < 0.05. (b) Niche networks for differentially enriched niches in HIV-(left) or HIV+ (right) samples. Nodes represent cell types and edge weights correspond to the number of unique samples with the corresponding interaction. Node size is proportional to connectedness, as measured by eigenvector centrality. (c) Comparison of the density of cells in the significant niches in the HIV+ samples broken down by p24 positivity. (d) Comparison of the proportion of CD8+ T cells that are positive for functional markers between significant niches in the HIV+ group, significant niches in the HIV-group, and all other niches. (e) Left: Heatmap showing the contribution of each cell type to the CD8+ T microenvironments (MEs). Right: Proportion of CD8+ T cells in each ME that are positive for functional markers. Values were column normalized. (f) Representative MIBI-TOF overlay and corresponding ME map, where colors correspond to the color bar in e (gray are cells that are not CD8+ T cells).

Network visualization of the niches discovered by QUICHE revealed that HIV-LNs exhibited diverse niches with many connections, as expected in a complex LN structure, whereas HIV+ LNs were less diverse and dominated by CD8+ T cells interacting with various myeloid and stromal cell types (Figure 3b). Investigating the interactions between CD8+ T cells and myeloid populations further, we found that the interactions of CD8+ T cells with CD14+ CD68+ CD163+ macrophages, CD14+ monocytes, CD11c+ DCs, and HLA-DR+ APCs were present in both HIV- and HIV+ LNs, and in HIV+ LNs, connections between CD8+ T cells and CD11c+ CD14+ monocytes were gained.

Stratifying the HIV+ population based on p24 positivity, we found that cells belonging to the significantly enriched niches in the HIV+ group, defined as having a spatial FDR less than 0.05 (Figure 3a), were present at higher frequencies in p24+ LNs compared to p24-LNs (Figure 3c). When we examined the two p24+ donors separately, we found that cells in these significant HIV+ niches were most abundant in the donor with ART interruption (Figure S6e). This suggests that viral reactivation of the HIV-1 reservoir following ART interruption is a primary driver of the observed changes in CD8+ T cell and myeloid interactions discovered using QUICHE.

Investigating the functional profile of the cells belonging to the differentially significant niches in LNs from the HIV+ group, we found that these cells expressed higher levels of activation markers such as CD45RO and Ki67 (Figure 3d). To further investigate caspase-1+ CD8+ T cells, which we found were elevated in p24+ LNs, we examined their spatial distribution. We found that these cells showed preferential clustering, being located closer to other caspase-1+ CD8+ T cells than to caspase-1-CD8+ T cells, suggesting potential feedback mechanisms that promote local activation (Figure S6f).

While QUICHE allowed us to find differentially significant spatial relationships, we found that many of the significant niches were highly enriched in the donor who had interrupted ART (Figure S6e). Furthermore, QUICHE downsamples the total number of cells for the analysis. To classify all CD8+ T cells based on their local neighborhoods and capture broad patterns of CD8+ T cell spatial organization, we identified CD8+ T cell microenvironments (MEs) by clustering CD8+ T cells according to their local neighborhood composition (Figure S6g-i). This analysis revealed four distinct MEs (Figure 3e-f). ME 1, characterized by CD8+ T cells surrounded by CD4+ T cells, including regulatory T cells (Tregs), was expectedly highest in HIV-LNs (Figure S6j). ME 1 was more enriched for CD8+ T cells expressing TCF1 and CD69 (Figure 3e). ME 2 CD8+ T cells were surrounded by other CD8+ T cells and were characterized by low levels of granzyme B and IFNγ, and high levels of CD45RO, PDL1, and galectin-9 (Figure 3e). This implies that aggregation of CD8+ T cells may modulate activation of adjacent T cells. We found that ME 2 was most abundant in LNs from p24+ tissues (Figure S6j), complementing our previous findings that p24+ donors displayed changes in CD8+ T cell activation and exhaustion. CD8+ T cells in ME 3 tended to be higher in both HIV+ groups and were located near various myeloid and stromal cells, often outside follicles. These cells expressed high levels of IFNγ, granzyme B, and Ki67, while having lower levels of TIGIT and caspase-1 than cells in ME 2. Finally, ME 4 CD8+ T cells were surrounded by B cells and FDCs, indicating that the CD8+ T cells infiltrating the follicle were in this cluster. This ME was most frequent in p24-LNs (Figure S6j).

Comparing the QUICHE niches with these CD8+ T cell MEs revealed that they capture complementary information (Figure S6k). As expected, the niche defined by CD8+ T cell-to-CD8+ T cell interactions consisted primarily of ME 2 cells, the ME characterized by CD8+ T cells surrounded by other CD8+ T cells. Interestingly, many other niches also contained cells from this ME, indicating that the MEs capture broader organizational patterns that complement the more specific cellular relationships identified by QUICHE.

Collectively, these analyses reveal that HIV-1 infection reshapes LN architecture, creating spatial niches dominated by clusters of CD8+ T cells with each other, in addition to CD8+ T cells in close association with myeloid and stromal populations. Such remodeling, particularly pronounced during viral activation, suggests that CD8+ T cells are recruited and activated at sites of persistent HIV-1 replication.

### CD56+ NK cells are depleted in HIV+ LNs

In the QUICHE spatial analysis, we found that many differentially expressed niches in the HIV-group involved CD56+ NK cells (Figure 3a-b). Given that NK cells are important effector immune cells involved in HIV-1 control and have been shown to respond rapidly in early infection^38,39^, we investigated NK cells in this cohort further. Although CD56+ NK cells composed a small percentage of all cells identified (0.86%), we found that they were depleted in HIV+ LNs, particularly in p24+ LNs (Figure S7a-c). Assessing the functional profile of these CD56+ NK cells, we found that Ki67 positivity was elevated in HIV-LNs and decreased in HIV+ LN. In contrast, there was higher detection of vimentin+ NK cells in p24+ LNs (Figure S7d), potentially contributing to functional suppression of these cells^40,41^. We also found that granzyme B trended higher in CD56+ NK cells from p24-LNs, while caspase-1 expression was lower. Investigating the co-expression patterns of granzyme B and caspase-1 further, we found that both markers had the highest co-expression with TIM3 and Ki67 (Figure S7e).

Assessing their localization in the tissue, we found that CD56+ NK cells were mostly located outside the follicles and displayed largely positive spatial enrichment scores with CD4+ T cells, though this enrichment score did not vary significantly between donor groups (Figure S7f-h). We also found a positive enrichment score with CD8+ T cells, which was slightly elevated in p24-LNs. In contrast, there was a decrease in the enrichment of CD11c+ CD68+ macrophages around CD56+ NK cells in p24+ LNs (Figure S7h). Taken together, these findings indicate that CD56+ NK cells are less frequent in tissue during viral activation and exhibit distinct functional states characterized by reduced proliferation and altered cytotoxic marker expression.

### Distinct immune microenvironments surround areas of active viral replication

Next, we asked how the immune microenvironment was remodeled around detected HIV-1 p24 signal, which reflects areas with active viral replication. To this end, we investigated the differences between individual follicles from the two PWH with strong p24 detection, one donor who was treatment-naïve and one donor with discontinued ART. This within-donor analysis allowed us to compare the characteristics of follicles in the same tissue while controlling for donor-specific factors. In the same LN, we found that some follicles showed strong p24 staining, while other follicles were negative for p24, demonstrating the heterogeneous nature of viral replication within LNs (Figure 4a, Figure S5a). Surprisingly, Tfh cells, which have previously been reported to be a major reservoir, showed no consistent trend between p24+ and p24-follicles, reflecting the complex role of these cells as reservoirs and in immune responses (Figure 4b). However, we did observe reduced numbers of Tregs and other CD4+ T cells in p24+ follicles (Figure 4b).

**Figure 4:**
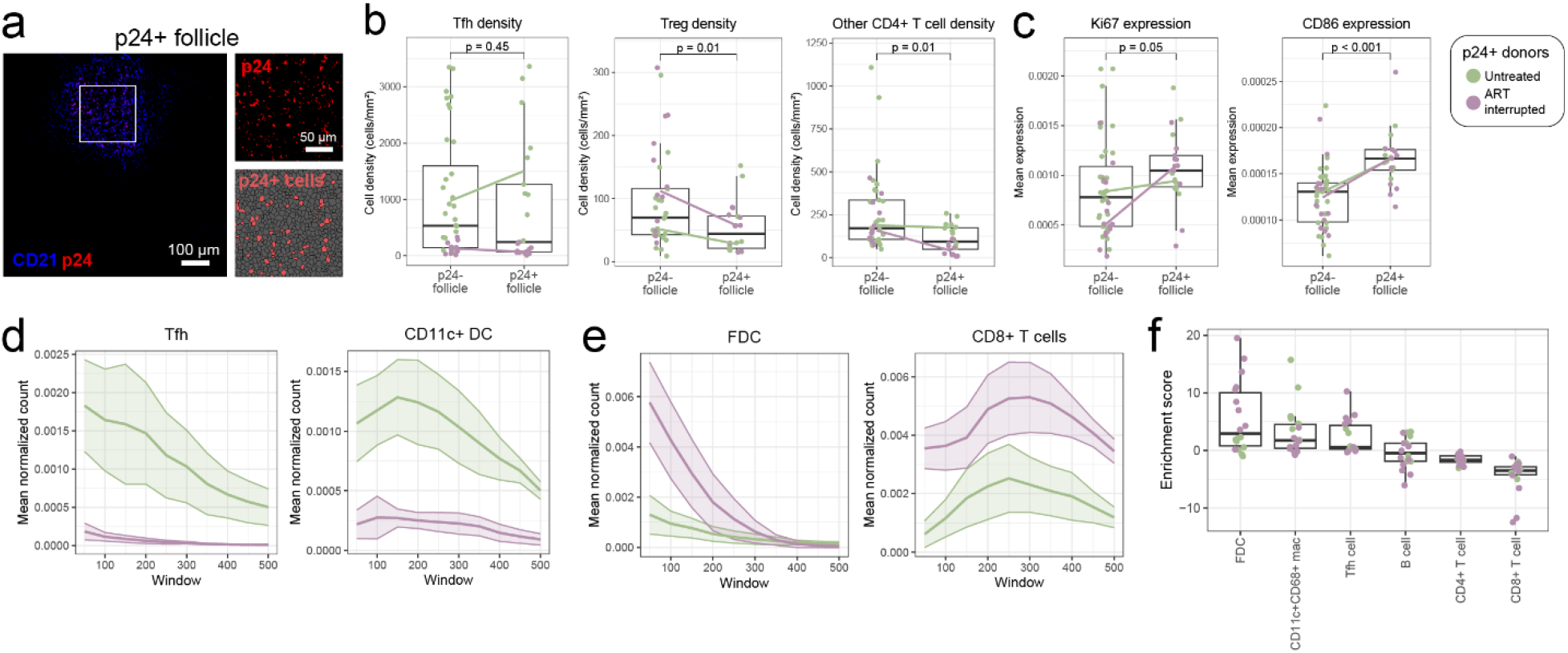
Active viral replication shapes distinct immune microenvironments in treatment-naive versus treatment-interrupted LNs. (a) Representative example of p24+ FOV. Top inset: p24 antibody staining. Bottom inset: p24+ cells in red, other cells in gray. (b) Comparison of Tfh cells, Tregs, and other CD4+ T cells in p24-vs p24+ follicles. Colors correspond to the two different p24+ donors. Each dot is one follicle. Lines connect the medians of the two donors. (c) Comparison of mean Ki67 and CD86 expression in p24-vs p24+ follicles. (d-e) Comparison of Tfh, CD11c+ DC, FDC, and CD8+ T cell density as a function of distance from p24+ cells. Lines represent the mean and shaded areas represent 95% confidence intervals. (f) Distribution of enrichment score between p24+ cells and other cell types. Each dot is one FOV. Enrichment scores were calculated by comparing the number of cell-cell contacts compared to a null distribution. To construct a null distribution, we use a bootstrapping approach by selectively randomizing the location of the cell populations of interest, calculating pairwise enrichment, and repeating this process 100 times.

Additionally, p24+ follicles exhibited higher mean expression of Ki67 and CD86, indicating increased immune activation (Figure 4c). We found that B cells expressed the highest levels of Ki67 (Figure S8a). CD8+ T cells also expressed higher levels of Ki67 in p24+ follicles compared to p24-follicles from the same LN (Figure S8b). We found that CD11c+ CD68+ macrophages had the highest expression of CD86, and various myeloid populations, including CD11c+ CD68+ macrophages, HLA-DR+ APCs, and CD11c+ DCs, expressed higher levels of CD86 in p24+ follicles (Figure S8c-d). These data indicate that there is localized activation in the myeloid compartment in response to viral replication in both the untreated and ART-interrupted settings, rather than having a systemic effect throughout the entire LN.

We then assessed the microenvironment around the detected p24 signal to assess how infection affects the local cellular neighborhood around the site of active viral replication (Figure S8e). Analysis of p24+ cells revealed distinct spatial associations depending on treatment status, suggesting that the cellular microenvironment around infected cells differs based on the host immune state and treatment history (Figure S8f-g). In the treatment-naïve donor, p24+ cells were more proximal to Tfh cells and CD11c+ myeloid cells (Figure 4d). In contrast, in the donor who had discontinued ART, p24+ cells were closer to FDCs and CD8+ T cells (Figure 4e).

We further performed an enrichment analysis, in which we determined which cell types were closest to the p24+ cells by comparing against a randomized null distribution to control for total cell numbers (Figure S6a). Surprisingly, enrichment analysis showed that CD8+ T cells were farther from p24+ cells than expected by random chance. This analysis supports the distance analysis shown in Figure 4e, which shows that the distance of CD8+ T cells to p24+ cells peaks at around 200-300 μm. Linking this enrichment analysis to our previous finding that caspase-1+ CD8+ T cells were associated with p24+ LNs, we assessed the enrichment of caspase-1- and caspase-1+ CD8+ T cells around p24+ signal. We found that caspase-1+ CD8+ T cells were significantly more enriched, though both enrichment scores were negative indicating farther distances than would be expected by random chance (Figure S8h). We found that CD8+ T cells expressing the checkpoint markers TIGIT and PD1 were closer to p24+ cells in the untreated sample but not in the setting of ART interruption, whereas we observed higher levels of GLUT1 in the setting of ART interruption, suggesting metabolic reprograming in CD8+ T cells from these tissues (Figure S8g). In contrast to the negative enrichment score of CD8+ T cells around p24+ cells, we found that p24+ cells had positive enrichment scores with FDCs, CD11c+CD68+ macrophages, and Tfh cells (Figure 4f). This spatial segregation indicates that despite increased CD8+ T cells in LNs from p24+ donors suggesting immune activation, these effector cells do not directly access sites of active viral reactivation, which tend to cluster near FDCs.

### Altered immune networks in LNs from aviremic PWH donors

Since most PWH achieve viremic suppression on ART, we specifically examined features most prominent in aviremic donors with undetectable plasma viral loads to understand the immunological landscape that characterizes successfully treated HIV-1 infection. We found that LNs from donors with suppressed plasma viremia contained higher numbers of B cells and fewer myeloid cells, including reduced CD11c+ dendritic cells, CD11c+CD14+ monocytes, and CD11c+CD68+ macrophages, when compared to tissue from PWH with detectable plasma viremia (Figure 5a). Interestingly, in these cell populations, LNs from viremic PWH more closely resembled LNs from PWOH. This shift in cellular composition suggests that viral suppression leads to a marked reduction in myeloid cell populations below even levels in HIV-LN, while B cell populations were elevated. Despite increased B cell numbers, these cells showed reduced proliferative activity (Figure S9a-b). Consistent with increased B cells, LNs from donors with suppressed plasma viremia showed the highest levels of CD8+ T cells in ME 4, which is composed of CD8+ T cells surrounded by B cells and FDCs (Figure S9c-d).

**Figure 5:**
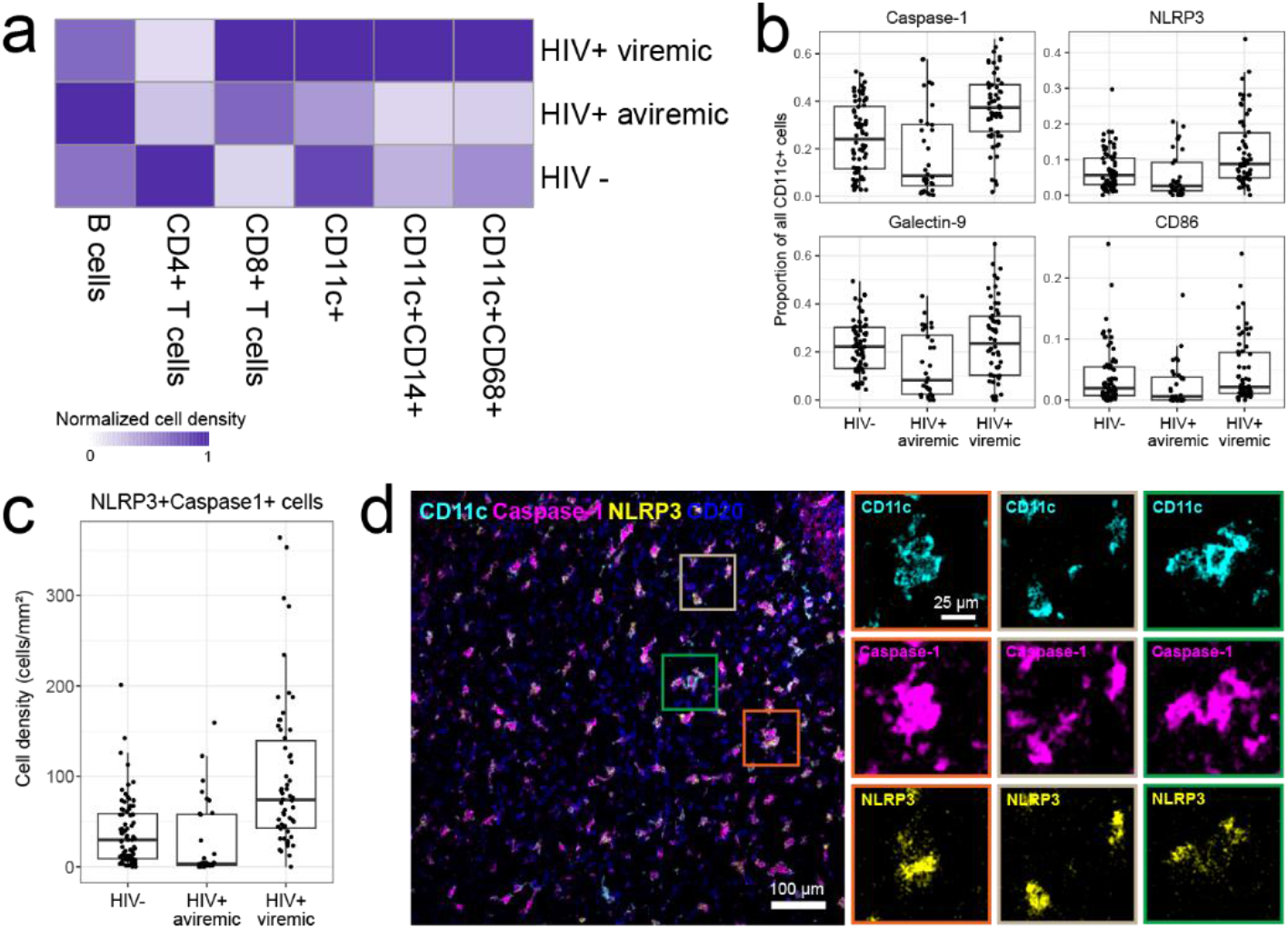
Viral suppression is associated with altered myeloid populations and reduced inflammasome activation. (a) Normalized mean cell density of various cell populations in viremic, aviremic, and HIV-donors. Aviremic donors are donors where the virus was undetectable in plasma. (b) Comparison of proportion of CD11c+ cells that are positive for functional markers. (c) Comparison of density of NLRP3+caspase-1+ cells. (d) Representative MIBI-TOF images.

Interestingly, the inflammasome markers caspase-1 and NLRP3 were lowest in the aviremic group and elevated in tissue from viremic individuals (Figure 5b-d). Investigating these markers further, out of all cells expressing either caspase-1 or NLRP3, only 4% of these cells co-expressed both markers (Figure S9e). The majority of inflammasome-positive cells were myeloid cells, and in particular cells that expressed CD11c (Figure S9f). Among all cells that expressed CD11c, the proportion expressing both NLRP3 and caspase-1 was reduced in the aviremic group, indicating reduced inflammasome activation in APCs during viral suppression (Figure S9g). Interestingly, we found a large proportion of cells that expressed caspase-1 and not NLRP3, suggesting a role for caspase-1 activation independent of NLRP3 (Figure S9h-i). This data shows that viral suppression leads to a dampening of inflammatory pathways in aviremic donors, even below HIV-levels, suggesting a potential association of the inflammasome with immune responses against the virus.

Altogether, this data shows that despite successful viral suppression in the plasma, the LN immune landscape is fundamentally altered, characterized by expanded B cell populations, reduced myeloid populations, and dampened inflammasome signaling.

## Discussion

In this study, we developed a spatial map of the cellular networks present in human LNs in HIV-1 infection, covering the spectrum of acute infection, successful viral suppression with ART, and viral reactivation due to ART interruption. Our findings reveal a paradoxical landscape of CD8+ T cell responses in HIV+ LNs. While we observed significant expansion of CD8+ T cells in HIV+ tissues, and to a greater extent in p24+ LNs, a substantial proportion of these cells lacked expression of canonical markers for activation, exhaustion, or effector function. This finding suggests that many CD8+ T cells may be recruited to LNs through non-specific inflammatory signals rather than antigen-driven activation. This interpretation is compatible with early reports of inflammatory signals such as MIP-1a mediating lymphocyte attraction to LNs^42^, although other studies reported an increase in proportions of memory CD45RO+ CD8+ T cells, rather than naive cells^43^. Supporting the trafficking of CD8+ T cells to LNs, we found that CD8+ T cells were preferentially enriched near stromal cells and endothelial cells, suggesting that tissue-derived chemotactic signals may drive their recruitment and retention. Importantly, this pattern persists even in people on suppressive ART with undetectable plasma viremia, suggesting that conserved inflammatory signals continue to drive CD8+ T cell accumulation during chronic HIV-1 infection despite systemic viral suppression.

In the last few years, accumulating evidence suggests that the ability of CD8+ T cells to infiltrate into B cell follicles while retaining cytotoxic markers and proliferative capacity may be associated with spontaneous control of HIV-1 infection^28,44^. In addition, studies in both primate and human tissue have shown that viral infection and ART initiation affect the recruitment of memory CD8+ T cells and expression of Th1 responses such as IFNγ, Tbet and the expression of tolerogenic markers such as Foxp3^29,45,46^. In our study, we expanded current knowledge to further characterize the phenotype of CD8+ T cells in follicular areas. Infiltrated cells closer to p24+ cells in follicular areas have been recently described to downregulate expression of granzyme B, potentially due to interaction of NKG2A with HLA-E in these areas^47^. In addition, previous studies have suggested that ligands for PD1 (PDL1) and TIGIT (PVR) are differentially expressed in HIV+ LNs and could participate in the modulation of CD8+ T cells in tissue^48^. Here, we have shown that inhibitory signals, including the ligands for checkpoint receptors such as galectin-9, are enriched in microenvironments characterized by CD8+ T cells interacting with other CD8+ T cells. Furthermore, while we have confirmed infiltration of CD8+ T cells into follicles, we found that they had a negative spatial enrichment score around p24+ cells. This suggests that despite overcoming anatomical barriers to follicular entry, CD8+ T cells may not efficiently engage infected cells at the sites of viral reactivation.

A key finding of our study is that despite similar follicular infiltration of CD8+ T cells in both p24- and p24+ LNs, the functional profile of these cells is different. CD8+ T cells inside the follicles in p24+ LNs were characterized by higher levels of caspase-1 and checkpoint markers PD1, TIGIT, and TIM3. Higher PD1 expression had previously been observed in tissue resident memory (Trm)-like CD8+ T cells expressing CD69 in tonsil tissue from HIV+ individuals^49^. In contrast, we found that CD8+ T cells displaying higher levels of LAG3, GLUT1, and IFNγ accumulated outside of follicles in p24+ LN. These findings are in line with previous studies that suggest that defective glycolysis and antiviral function are associated with differential patterns of TIM3 versus TIGIT expression on CD8+ T cells from PWH^19^. In addition, higher TIGIT and LAG3 expression have been described to be more enriched in CD8+ T cells in tissue from HIV-1 chronic progressors compared to elite controller individuals^44^.

In addition to the functional phenotype of CD8+ T cells, our study addresses the relationships with innate pathways that may be relevant to chronic inflammation in HIV+ LNs^50–52^. We observed an enrichment in myeloid cells expressing inflammasome NLRP3 and caspase-1 in LNs from viremic individuals, which is consistent with previous studies that have suggested that inflammasome activation in macrophages contributes to non-AIDS inflammatory disorders^53^. We found that caspase-1 expression is enriched in CD11c+ cells, suggesting that DCs may play a key role modulating inflammation in tissue. In addition, our data suggests that a proportion of caspase-1+ cells are not from myeloid origin, particularly CD8+ T cells. In this regard, NLRP3 inflammasome has been shown to potentially regulate tissue-resident CD8+ T cells in lupus nephritis, and its role in HIV-1 infection warrants further investigation^54^. The elevated expression of caspase-1 independent of NLRP3 in CD8+ T cells from p24+ tissues suggests potential alternative inflammasome pathways that may affect the function of these cells. In addition to NLRP3, increased expression of alternative inflammasome sensors induced by viral replication in CD4+ T cells has been reported in PWH^55^, but their implication in CD8+ T cells remains to be explored. Conversely, the marked reduction in inflammasome signaling in aviremic donors, even below levels observed in HIV-controls, indicates that successful viral suppression may be achieved in part by decreasing the activity of myeloid cells. Notably, our study included a viral protein marker. However, the HIV-1 reservoir is heterogeneous and some infected cells, particularly in tissue, may express viral mRNA that is not translated into protein^7^. Sensing of HIV-1 mRNA products has been associated with activation of inflammasomes other than NLRP3 in myeloid cells^56^. Therefore, differences in myeloid distribution and inflammasome activation observed in p24-LN that differ from PWOH could be attributed to viral RNA sensing and further studies including detection of HIV-1 mRNA would be needed to validate this hypothesis.

In our study, we found that tissue p24 signal was located near FDCs in germinal centers, in agreement with previous studies^29,57,58^. Our findings suggest that FDCs may contribute to the latent reservoir that is being reactivated in the setting of ART interruption, while Tfh cells may account for viral spreading in acute infection. Our observation that viral activation, particularly following ART interruption, occurs preferentially in proximity to FDCs has important implications for understanding HIV-1 persistence. FDCs are known to trap and present antigens for extended periods^58–60^, and their location within germinal centers may provide an immunologically privileged microenvironment that supports viral persistence. However, these findings are based on analysis of tissue from two donors and therefore additional validation in larger cohorts with similar treatment histories is required.

An additional aspect of this study that has not been explored is how the phenotype and distribution of other immune cells not targeted in our panel may be relevant, such as CD56-NK cells, and how they may vary with active or suppressed viral replication. Recent studies suggest that expression of HLA-E is associated with dysfunctional CD8+ T cells in HIV+ LN^47^. Taking into consideration that this molecule is a ligand of the inhibitory NKG2A and activating NKG2C receptors expressed by NK cells, additional studies should be conducted to further address changes in specific NK cell subsets in human HIV+ LN.

An important caveat from the study is that all donors, both HIV+ and HIV-, had lymphadenopathies that were suspected for cancer. Lymph nodes are rarely removed from healthy donors, and we tried to select for pathologies that would result in minimal immune alterations (no metastatic cells were found in any of the analyzed samples). However, the immune landscape of these donors may be influenced by systemic immune responses, and therefore additional validation studies performed in tissue from donors who do not have pathologies associated with immune activation should be performed.

These findings have important implications for HIV-1 cure strategies. The persistence of altered immune architecture even during successful viral suppression suggests that tissue-level interventions may be necessary to restore normal immune function. The spatial segregation of CD8+ T cells from sites of viral activation highlights potential limitations of current immunotherapeutic approaches and suggests that strategies to enhance CD8+ T cell trafficking to, or retention at, sites of viral persistence may be necessary for effective reservoir clearance.

## Methods

### Tissue cohort

We obtained LN tissue from five PWH and five PWOH from the Pathology Anatomy Unit at Hospital Universitario La Princesa (Madrid, Spain) in collaboration with Dr. Magdalena Adrados de Llano. The tissues used in the study were obtained from paraffin-fixed tissue sample blocks previously obtained for clinical analyses under an informed consent (Register No. 3518) approved by Hospital La Princesa’s Bioethical committee. All selected LN blocks had no metastatic infiltration. We matched sex and age between PWH and PWOH. Further clinical characteristics of the different samples can be found in Supplementary Table 1.

### Panel construction

Most antibodies used in this study were previously validated for MIBI-TOF^61–65^. Newly selected target antibodies used for this study were validated via immunohistochemistry to confirm appropriate specific staining in control tissues (Figure S2a). All antibodies were metal-labeled using the Ionpath conjugation kit according to the manufacturer’s protocol. Calprotectin and mast cell tryptase were conjugated to Ga69 and Ga71 respectively using monomeric maleimido-mono-amide NOTA. To prevent degradation and prolong shelf life, labeled antibodies were lyophilized individually with 100 mM trehalose in 1 µg or 5 µg aliquots. Titer optimization for new targets was performed using serial dilutions (1 µg/mL, 0.5 µg/mL, 0.25 µg/mL, and 0.125 µg/mL) or the recommended titer for previously validated MIBI-TOF antibodies. The full panel can be found in Supplementary Table 2.

### MIBI-TOF staining

Tissues were sectioned (5 μm section thickness) from tissue blocks on gold-sputtered slides. Slides were baked at 70°C overnight, followed by deparaffinization and rehydration with washes in xylene (3x), 100% ethanol (2x), 95% ethanol (2x), 80% ethanol (1x), 70% ethanol (1x) and ddH2O with a Leica ST4020 Linear Stainer (Leica Biosystems). Tissues next underwent antigen retrieval by submerging sides in 3-in-1 Target Retrieval Solution (pH 9, DAKO Agilent) and incubating at 97°C for 40 min in a Lab Vision PT Module (Thermo Fisher Scientific). After cooling to room temperature, slides were washed in 1x PBS IHC Washer Buffer with Tween 20 (Cell Marque) with 0.1% (w/v) bovine serum albumin (Thermo Fisher). Next, all tissues underwent two rounds of blocking, the first to block endogenous biotin and avidin with an Avidin/Biotin Blocking Kit (BioLegend). Tissues were then mounted onto a Sequenza system, washed with wash buffer, and blocked for 1 h at room temperature with 1× TBS IHC Wash Buffer with Tween 20 with 3% (v/v) normal donkey serum (Sigma-Aldrich), 0.1% (v/v) cold fish skin gelatin (Sigma-Aldrich), 0.1% (v/v) Triton X-100, and 0.05% (v/v) Sodium Azide. The first antibody cocktail was prepared in 1x TBS IHC Wash Buffer with Tween 20 with 3% (v/v) normal donkey serum (Sigma-Aldrich) and filtered through a 0.1-μm centrifugal filter (Millipore) prior to incubation with tissue overnight at 4 °C in the Sequenza. Following the overnight incubation, slides were washed twice for 5 min in wash buffer. The second day, an antibody cocktail was prepared as described and incubated with the tissues for 1 h at 4°C. Following staining, slides were washed twice for 5 min in wash buffer and fixed in a solution of 2% glutaraldehyde (Electron Microscopy Sciences) solution in low-barium PBS for 5 min. Slides were washed in PBS (1x), 0.1 M Tris at pH 8.5 (3x) and ddH2O (2x) and then dehydrated by washing in 70% ethanol (1x), 80% ethanol (1x), 95% ethanol (2x) and 100% ethanol (2x). Slides were dried under vacuum prior to imaging.

Detailed protocols can be found at the links below:

Reagent preparation:dx.doi.org/10.17504/protocols.io.bhmej43e

IHC staining:dx.doi.org/10.17504/protocols.io.bf6ajrae

MIBI staining:dx.doi.org/10.17504/protocols.io.dm6gprk2dvzp/v5

Sequenza staining:dx.doi.org/10.17504/protocols.io.bmc6k2ze

Antibody lyophilization:dx.doi.org/10.17504/protocols.io.bhmgj43w

Lyophilized antibody reconstitution:dx.doi.org/10.17504/protocols.io.bjd6ki9e

### MIBI-TOF imaging

All MIBI data was generated on a commercial MIBIScope instrument (IonPath, Menlo Park, USA). For each LN tissue, we used paired IHC or H&E images to identify the fields-of-view (FOVs) that would be acquired on MIBI, sampling from the various regions of the LN (Figure S1b). Prior to imaging, the slides were sputtered with 10 nm of gold to help with uniform imaging. We used the same settings across all MIBI data acquired in this study. To pre-raster the gold coat before image acquisition, we used a setting of 40 nA and a dwell time of 0.25 ms for each FOV. For image acquisition, the size of each FOV was 800 μm x 800 μm / 2048 pixels x 2048 pixels. We used the “Coarse” settings on the MIBIScope instrument. Between all runs, we conducted an imaging test on the molybdenum and PMMA standards per the manufacturer’s recommendations and settings. We used the autogain feature on the MIBIScope instrument to automatically adjust the detector gain during a run. To minimize image artifacts from instrument drift, we randomized the acquisition order of the FOVs. All default image processing (background correction and denoising) were disabled.

### Image compensation with Rosetta

Low-level processing of the MIBI-TOF images was conducted with our publicly available, in-house pipeline, Toffy (https://github.com/angelolab/toffy). Following image extraction, we removed sources of signal contamination using a compensation method called Rosetta. This approach implements channel-by-channel and pixel-by-pixel image compensation based on defined rates of signal contamination provided from the manufacturer (IonPath, Menlo Park, CA) and experimentally determined. The coefficients we used for subtraction were determined manually by image inspection. The source and target channels, as well as the coefficients used for subtraction can be found in Supplementary Table 3.

### Cell segmentation and classification

We used Mesmer to segment all images in the cohort^66^. Mesmer is a pre-trained deep learning model that takes two channels of input data, a nuclear channel and a membrane channel. We used the HH3/dsDNA marker as the nuclear channel, and combined the HLA-I, CD45, and CD31 markers as the membrane channel. We then ran Mesmer using the default parameters as part of our publicly available, in-house pipeline, ark-analysis (https://github.com/angelolab/ark-analysis). Representative images of the resulting segmentation can be found in Figure S3a.

Using ark-analysis (v0.8.0), we performed pixel clustering using Pixie, a clustering algorithm optimized for multiplexed image data^67^. We used the following parameters: blur_factor = 2 and subset_proportion = 0.1. We used these phenotypic markers for clustering: Calprotectin, CD3e, CD4, CD8a, CD11c, CD14, CD20, CD21, CD31, CD45, CD56, CD68, CD163, CXCR5, FOXP3, HLADR, MastCellTryptase, PD1, SMA. We initially over-clustered the data into 200 clusters which were then combined into 41 meta-clusters using consensus hierarchical clustering and manual adjustments by examining the heatmap of marker expression in each cluster. We next determined the proportion of each pixel cluster in each segmented cell object. These profiles were then used to cluster cells, first over clustered into 200 clusters and then combined into 20 metaclusters using consensus hierarchical clustering and manual adjustments. To confirm accurate clustering and correct systemic instances of inaccurate assignment, we inspected the clusters in Mantis Viewer^68^.

We used Nimbus to refine cell type assignments and determine functional marker expression status^69^. Nimbus is a deep learning model that predicts marker positivity from multiplexed imaging data. Nimbus generates a confidence score for each marker in every cell, indicating the likelihood of marker expression in that cell. Using these Nimbus scores, we reassigned cells in the “Other” and “Unassigned” clusters that had high scores for lineage markers. These thresholds were determined empirically by inspecting the images and Nimbus scores. To confirm accurate clustering and correct systemic instances of inaccurate assignment, we inspected the clusters in Mantis Viewer. We then manually evaluated Nimbus scores to establish marker-specific thresholds for classifying cells as positive or negative for each functional marker. These thresholds can be found in Supplementary Table 4.

### Lymph node region masks

To create masks of the follicular and extrafollicular areas, we used a combination of the MIBI-TOF images of CD21 and Ki67. We Gaussian blurred the images using a large sigma (100) and used Otsu thresholding to binarize the images. These images were then manually adjusted in ImageJ and combined to create follicular masks. Extrafollicular masks were created by subtracting the follicular mask from the total tissue area. Because MIBI-TOF is performed on gold slides, we used the signal from the gold channel that is acquired by the instrument to identify regions of empty slide. By subtracting out regions of empty slide from each image, we calculated the total tissue area in each image.

### Feature extraction pipeline and mixed-effect model

To systematically compare HIV+ and HIV-LNs, we extracted a total of 1189 features for each image. These features included cell density, cell ratios, functional marker expression, cellular microenvironments, cell-cell enrichment scores, and neighborhood diversity. For cell density, we counted the number of each cell type in each image, then divided the count by total tissue area in the image^70^. We also performed this cell density calculation for each cell population specifically in the follicular or extrafollicular regions using the masks generated as described above. For cell ratios, we calculated the ratio of various cell populations, including CD4+ T cells to CD8+ T cells, Tregs to total CD4+ T cells, and total T cells to total myeloid cells. For functional marker expression, we determined thresholds for positivity using the Nimbus scores by manual inspection. Using these thresholds, we calculated the proportion of each cell type that was positive for each functional marker, as well as the cell density of each functional marker-positive cell type. We also calculated the mean expression of each of the functional markers in each cell population for every image. For cellular microenvironments, we used the Spatial-LDA algorithm, an adaptation of the topic modeling method latent Dirichlet allocation for finding spatial patterns of cells in tissue^71^. Spatial-LDA was implemented as described previously, with spatial radius r = 100 μm, and microenvironment number of n = 6. The microenvironment number was determined empirically using the metrics inertia and Silhouette score^62,71^. For the cell-cell enrichment scores, we used an approach as previously described, in which we compared the number of observed cell-cell contacts with a randomized null distribution^63^. For every image, for each pair of cell populations X and Y, the number of times Y cells within a 20 μm radius of X cells was counted. A null distribution was produced by performing 100 bootstrap permutations where the location of Y cells was randomized in the image and the number of X-Y contacts was counted in the randomized image. The number of true X-Y contacts was then compared against this null distribution to generate a z-score. For neighborhood diversity, for each cell, we counted the number of each cell type in a radius of 50 pixels around that cell, then computed the Shannon Diversity Index, a popular metric in ecology, to quantify the heterogeneity of the local cellular microenvironment. Then we found the average Shannon Diversity Index for each cell population in each image.

To test for differences between HIV+ and HIV-LNs, we used a mixed-effect model to control for multiple images from the same LN. The fixed effect was HIV+ or HIV-, while the random effect was the donor to account for person-to-person variability. We used the nlme package in R. We also calculated Cohen’s d, the standardized effect size, for each feature. P-values were generated using the mixed-effect model.

### QUantitative InterCellular nicHe Enrichment (QUICHE) spatial analysis

To test for differences in cellular organization across conditions (i.e. HIV-vs HIV+ LNs), we performed spatial statistical differential abundance testing using QUICHE (https://github.com/jranek/quiche)^37^. Briefly, we identified local cellular niches within each image according to spatial proximity by enumerating the number of each cell type within a set radius (radius = 200 pixels), performed distribution-focused downsampling to select a subset of niches (niches = 50,000) from each sample for condition comparisons, and constructed a niche similarity graph for differential enrichment testing (n_neighbors = 30, min_cell_threhsold = 3, k_sim = 100).

QUICHE niche neighborhoods were considered significant if spatial false discovery rate (FDR) < 0.05.

### CD8+ T microenvironment analysis

To classify all CD8+ T cells based on their local neighborhood, for each CD8+ T cell, we enumerated the number of each cell type within a 50 pixel radius. Using these counts as the feature vector for each CD8+ T cell, we used k-means clustering to cluster the cells into four microenvironment clusters. The number of clusters was determined empirically using inertia and Silhouette score, and by manual inspection of cluster heatmaps.

### p24+ follicle analysis

From the two PWH donors that showed positive p24 staining in any imaged area of the LN, we classified each follicle in all images from these donors as p24+ or p24-based on detectable p24 staining within the follicle. We determined the threshold for p24 positivity using Nimbus scores and manual inspection of images. We then assessed the differences between the p24+ and p24-follicles.

To assess the neighborhood around p24+ signal as a function of distance, we counted the number of each cell type in 50 μm increments around the center cell that showed p24 positivity, from 0 μm to 500 μm (i.e. 0 μm - 50 μm, 50 μm - 100 μm, etc.). These counts were then normalized by the area of each window. We also found the proportion of CD8+ T cells that were expressing functional markers in each of these windows.

For the enrichment analysis, we used the same approach as described above^63^, in which we compared the number of observed contacts between p24+ signal and all cell types with a randomized null distribution. For every image, for each cell population X, the number of times X cells within a 20 μm radius of p24+ signal was counted. A null distribution was produced by performing 100 bootstrap permutations where the location of X cells was randomized in the image and the number of contacts was counted in the randomized image. The number of true contacts was then compared against this null distribution to generate a z-score. For FDCs, cells were only randomized within the follicle mask.

## Figure generation

Figures were produced using Adobe Illustrator, Adobe Photoshop, and BioRender.

## Data availability

Imaging data and metadata are deposited in the BioImage Archive at https://doi.org/10.6019/S-BIAD2864.

## Code availability

Code used to perform analyses and generate figures are available at https://github.com/angelolab/publications/tree/main/2026-Liu_Calvet-Mirabent_etal_HIV-LN.

## Acknowledgements

C.C.L. was supported by grant no. NIAID F31AI165180 and the Stanford Graduate Fellowship. M.C.M. was supported by La Caixa Banking Foundation ETI-CureHIV (HR20-00218), Boehringer Ingelheim Fonds Travel grant, Asociación Red de Investigación En Sida (RIS) and CFAR Mentored Scientist Award (CA-0259646). M.A. was supported by NIH grant nos. 5U54CA20997105, 1R01CA24063801A1, 5R01AG06827902, 5UH3CA24663303, 5R01CA22952904, 1U24CA22430901, 5R01AG05791504, 5R01AG05628705, 5U24CA22430903, 3U54HL165445-03S1, 5R01AG05628705, 5R01AG05791505; the Department of Defense grant nos. W81XWH2110143, 5U54CA261719-05. E.M.G was supported by the Spanish Agencia Estatal de Investigación RETOS, Generación de conocimiento and consolidation programs (PID2021-127899OB-I00; CNS2023-144841; PID2024-160973OB-I00), GLD24/00117 grant from Gilead biosciences, La Caixa Banking Foundation ETI-CureHIV (HR20-00218) and infectious diseases CIBER from ISCIII (CB21/13/00107).

## Declaration of interests

M.A. and S.C.B. are inventors on patents related to MIBI technology (patent nos. US20150287578A1, WO2016153819A1 and US20180024111A1). E.M.G. declares no competing interests.

## Author contributions

M.C.M. and E.M.G. conceived the project and provided tissue samples. M.C.M, M.A.L., and E.M.G. curated the tissue specimens and associated metadata. C.C.L., M.C.M., and E.M.G. designed the MIBI-TOF panel. C.C.L., M.C.M., A.S., and M.G. performed reagent optimization and generated the MIBI-TOF data. C.C.L. processed and analyzed the MIBI-TOF data. C.C.L., M.C.M., and E.M.G. wrote the manuscript. S.C.B., M.A., and E.M.G. supervised the project. All authors provided feedback on the manuscript.

