## Supplementary Figures for "Spatial proteomics reveals CD8+ T cell signatures and cellular niches associated with active HIV-1 replication in lymph nodes"

Supplementary Figure 1

a

HIV+ samples

| HIV positive | On ART | Virus detectable in plasma | p24 detectable in LN | Acute | ART interrupted |
|---|---|---|---|---|---|
| ✓ | ✓ | | | | |
| ✓ | ✓ | | | | |
| ✓ | | ✓ | | | |
| ✓ | | ✓ | ✓ | ✓ | |
| ✓ | | ✓ | ✓ | | ✓ |

b FOV selection for MIBI-TOF

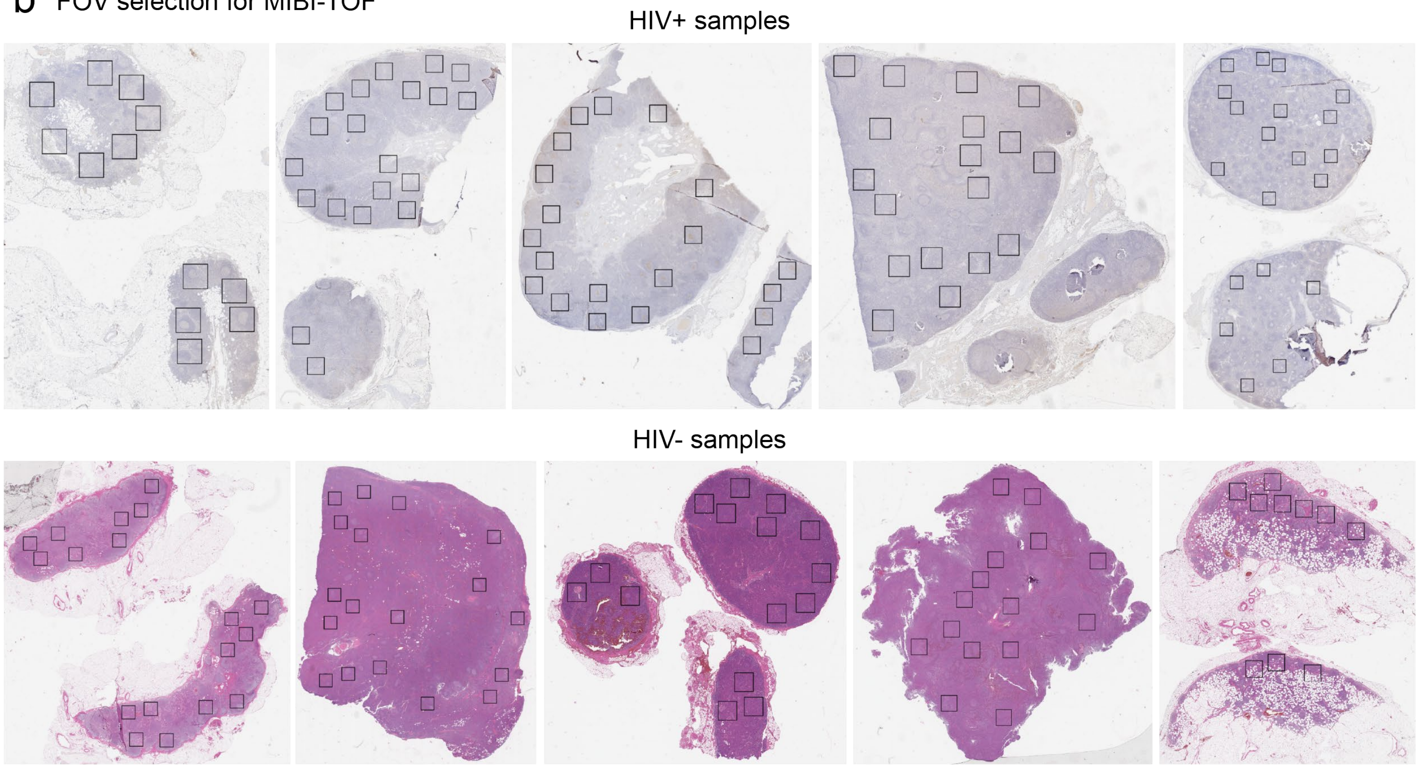


**Supplementary Figure 1: Sample details and region selection for MIBI-TOF imaging**

(a) Cohort details of HIV+ samples. Each row is one sample. (b) Region selection for MIBI-TOF imaging using p24 chromogenic immunohistochemistry stains (HIV+ samples) and H&E images (HIV- samples). Each square is 1 FOV (800 µm x 800 µm).

### Supplementary Figure 2

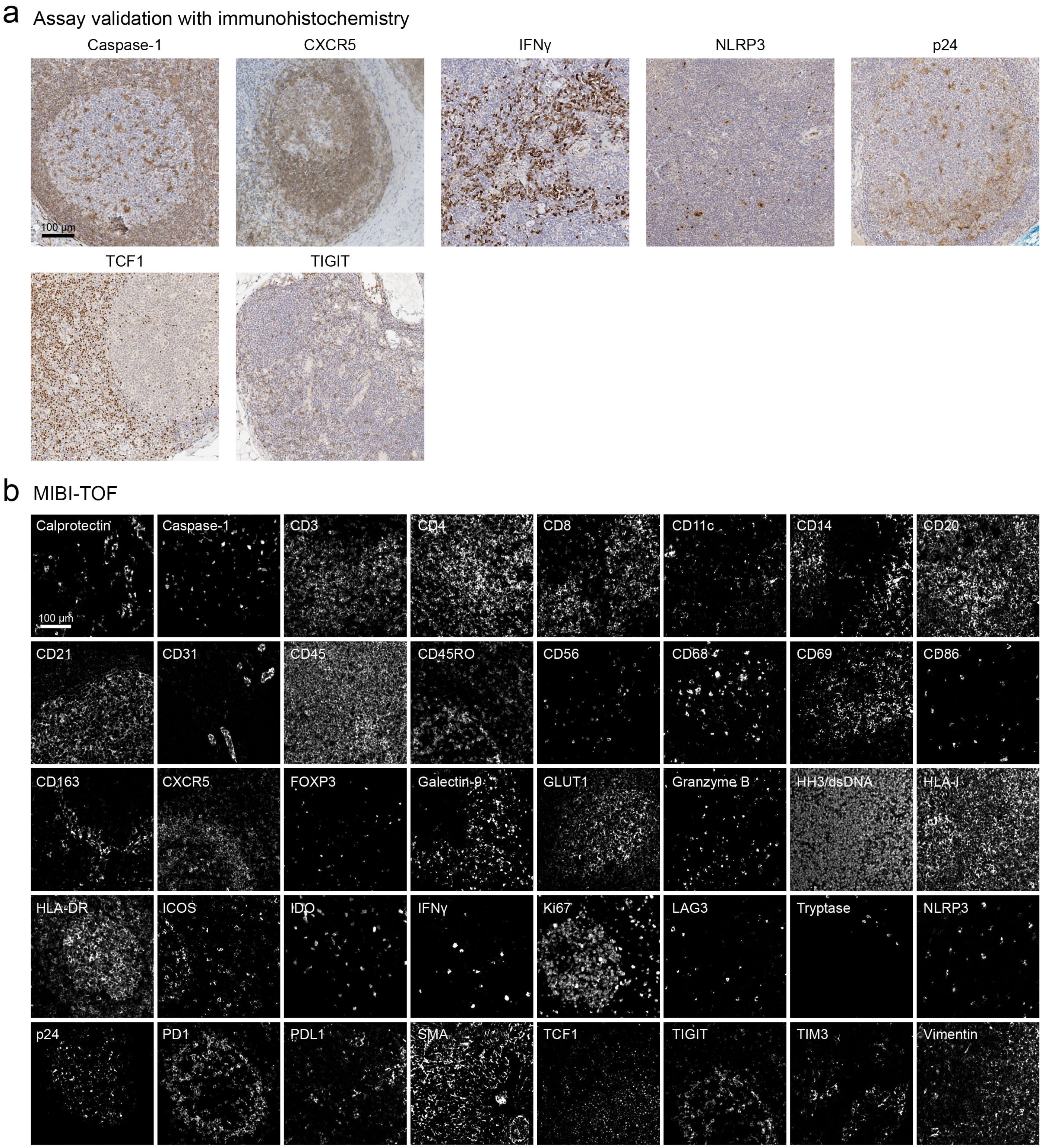


**Supplementary Figure 2: Assay validation**

(a) Representative chromogenic immunohistochemistry images for newly validated antibody targets for this study in lymph node tissue. Validation for other antibody targets in the panel can be found in previous publications.[61-65] (b) Representative MIBI-TOF images for all antibody targets in lymph node tissue.

### Supplementary Figure 3

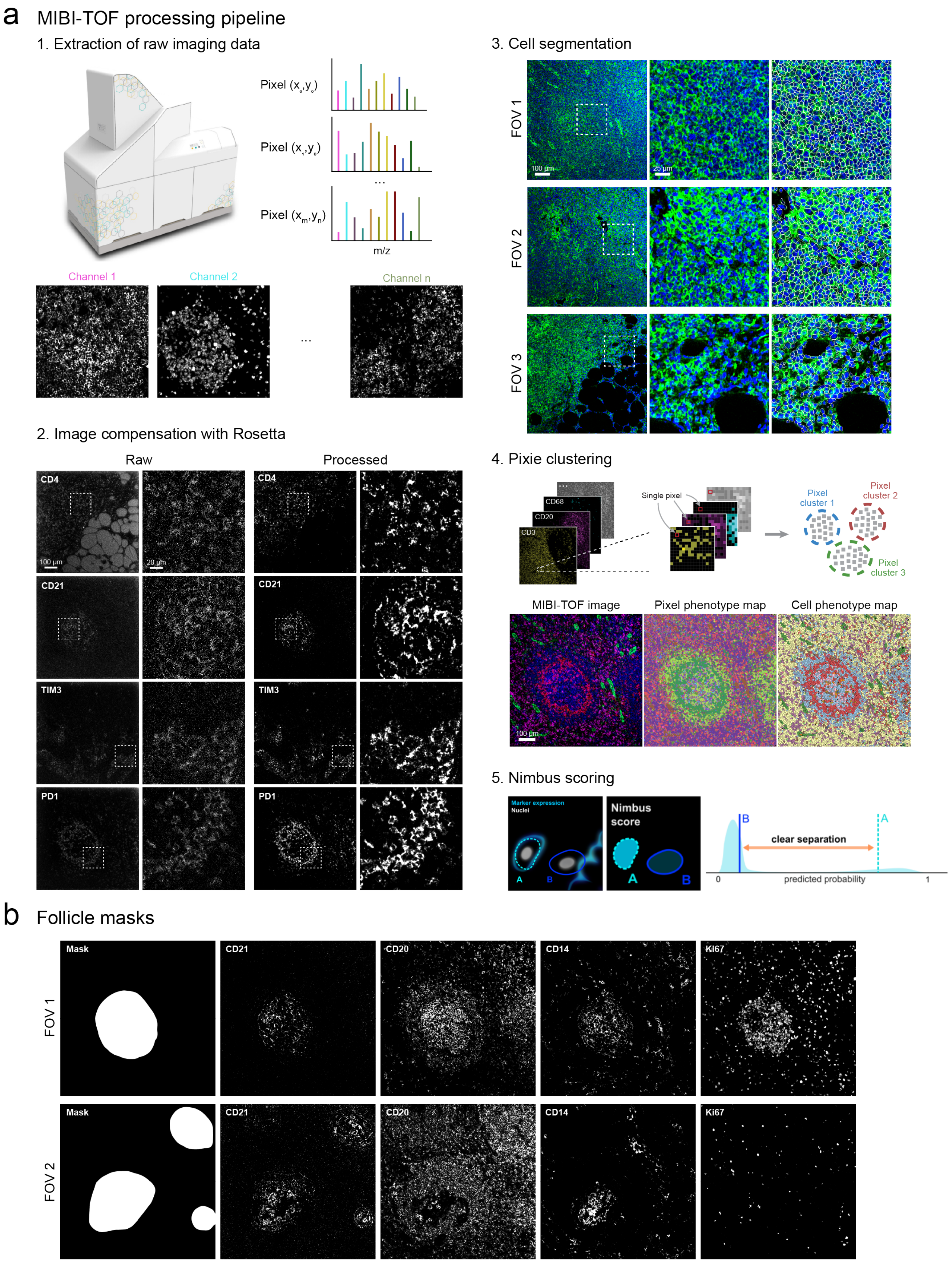

**Supplementary Figure 3: Computational pipeline for MIBIT-TOF image analysis**

(a) Overview of MIBI-TOF processing pipeline. After image extraction, images were compensated using Rosetta (see Methods). Cells were then segmented using the Mesmer algorithm. Blue is the nuclear marker (HH3/dsDNA) and green is the membrane marker (HLA-I, CD45, and CD31). Cells were then classified using Pixie and Nimbus. (b) Example follicle masks.

Supplementary Figure 4

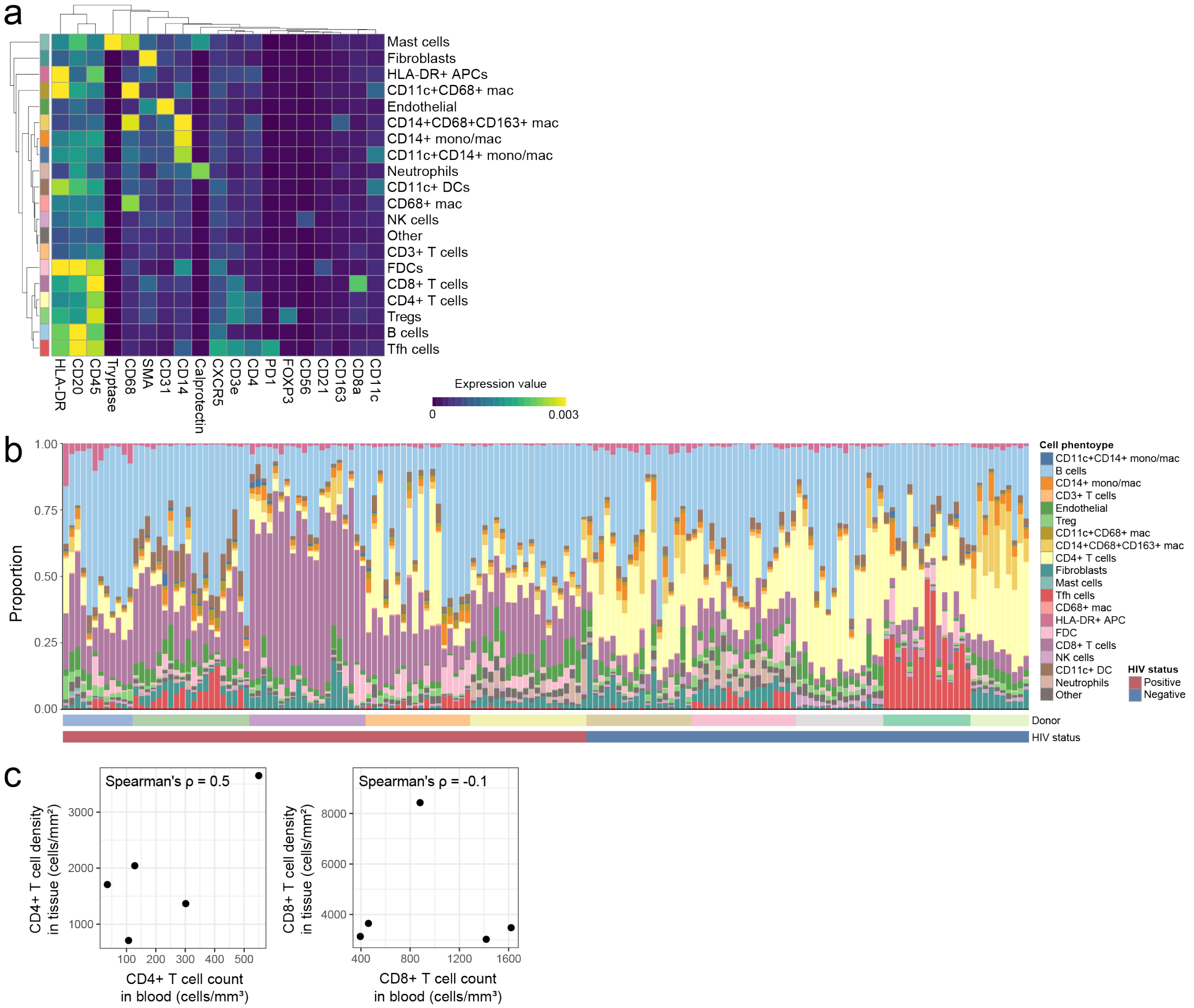


**Supplementary Figure 4: Cell type identification and quantification of MIBI-TOF images**

(a) Heatmap of cell phenotypes without column z-score normalization. Values are mean expression values from the processed images. (b) Frequency of the identified cell phenotypes for each FOV in the study. (c) Correlation of CD4+ T cell and CD8+ T cell counts in the tissue and blood at the time of tissue sampling.

Supplementary Figure 5

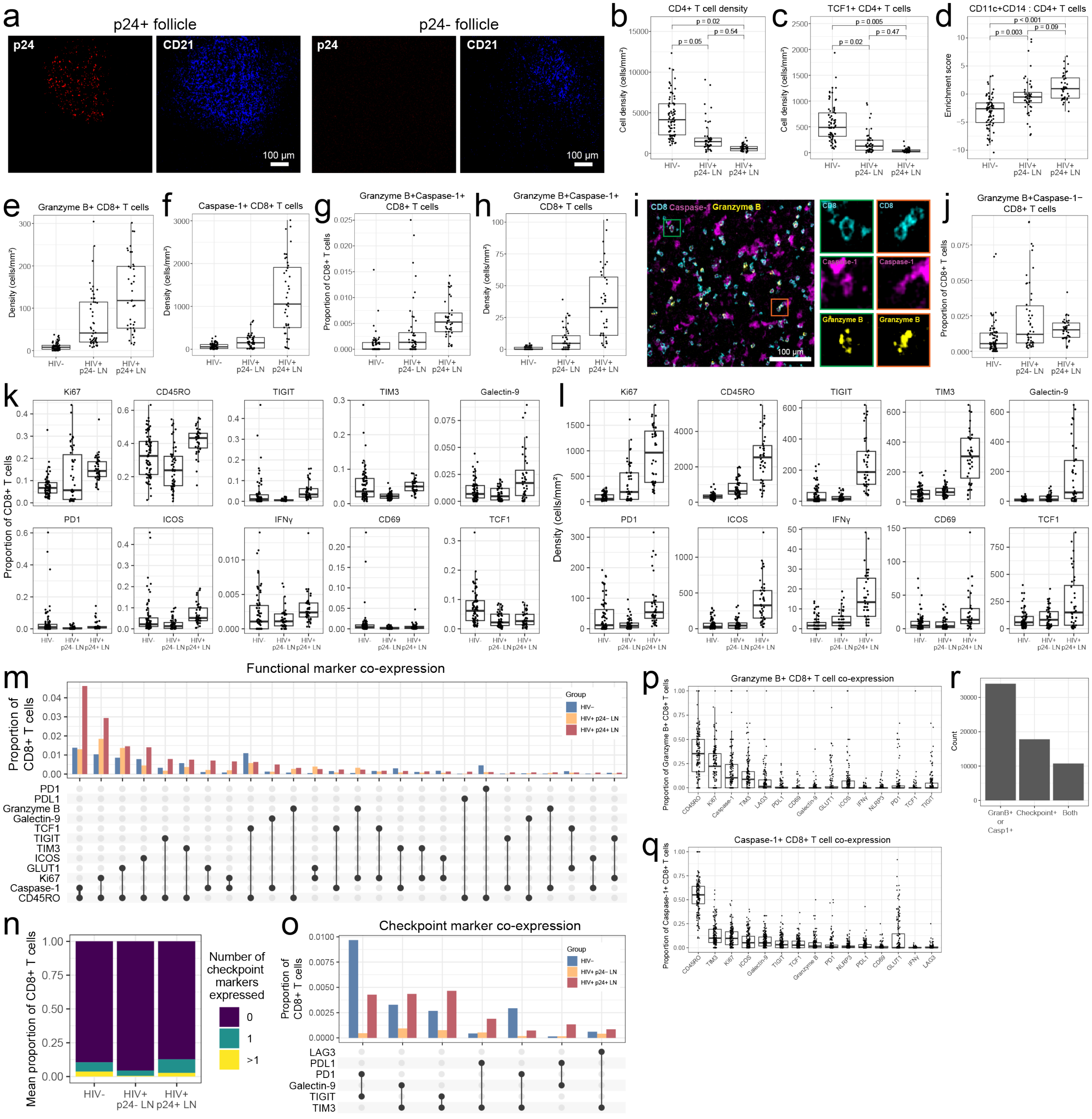

**Supplementary Figure 5: Phenotypic profiling of T cells in LN from PWH and PWOH**

(a) Representative examples of p24 tissue staining in two follicles from the same LN, one follicle that is p24+ and the other is p24-. (b) Comparison of CD4+ T cell density between donor groups. (c-d) Using a mixed-effect model to determine significance, the most significantly different features in the HIV- (c) and HIV+ (d) groups. (e) Comparison of granzyme B+ CD8+ T cell density. (f) Comparison of caspase-1+ CD8+ T cell density. (g) Comparison of proportion of CD8+ T cells that are positive for granzyme B and caspase-1. (h) Comparison of density of granzyme B+caspase-1+ CD8+ T cells. (i) Representative MIBI-TOF image of granzyme B+caspase-1+ CD8+ T cells. (j) Comparison of proportion of CD8+ T cells that are positive for granzyme B and negative for caspase-1. (k) Proportion of functional marker positive CD8+ T cells and (l) functional marker cell density. (m) Co-expression patterns in CD8+ T cells in different donor groups. Co-expression partners with at least 100 cells in p24+ LNs are shown. (n) Proportion CD8+ T cells that express checkpoint markers. (o) Co-expression patterns of checkpoint markers in CD8+ T cells. Co-expression partners with at least 100 cells are shown. (p) Proportion of granzyme B+ or (q) caspase-1+ CD8+ T cells that express other functional markers. (r) Number of CD8+ T cells that express granzyme B or caspase-1, any checkpoint marker, or both.

Supplementary Figure 6

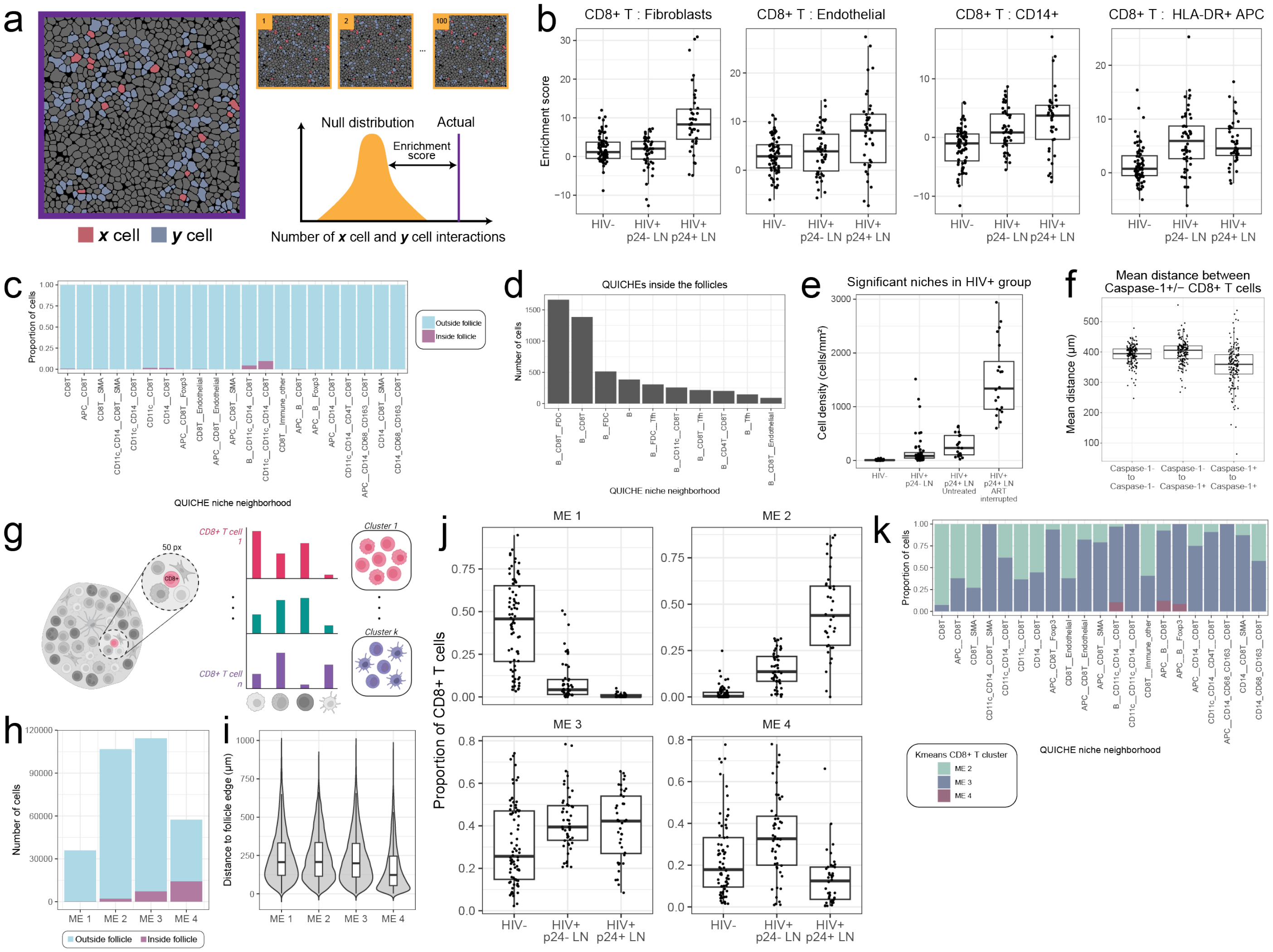


**Supplementary Figure 6. QUICHE and CD8+ T cell microenvironment analysis**

(a) Conceptual overview of cell-cell enrichment score calculation. Enrichment scores were calculated by comparing the observed number of cell-cell contacts compared to a null distribution generated by bootstrapping (see Methods). (b) Comparison of cell-cell enrichment scores for CD8+ T cells. (c) Proportion of cells that are inside or outside the follicles for the significant differentially enriched QUICHE niches in the HIV+ group (median logFC > 1 and median Spatial FDR < 0.05). (d) Breakdown of the most frequent QUICHE niches inside follicles (top 10). (e) Comparison of significant niches in the HIV+ group, where the p24+ donors were separated by treatment status. (f) The mean distance between caspase-1+ and caspase-1- CD8+ T cells. (g) Conceptual overview of CD8+ T cell microenvironment (ME) analysis. For every CD8+ T cell in the image, a 50 pixel (~20 µm) radius was used to determine the frequency of cells in its local neighborhood. All CD8+ T cells were then clustered using k-means clustering into 4 MEs. (h) Number of cells that are inside or outside the follicle for the ME clusters. (i) Distance of the CD8+ T cells in each ME cluster to the follicle edge. (j)

Comparison of the density of each ME cluster across donor groups. (k) Proportion of cells in each significant differentially enriched QUICHE niche that belong to each CD8+ T cell ME cluster.

#### Supplementary Figure 7

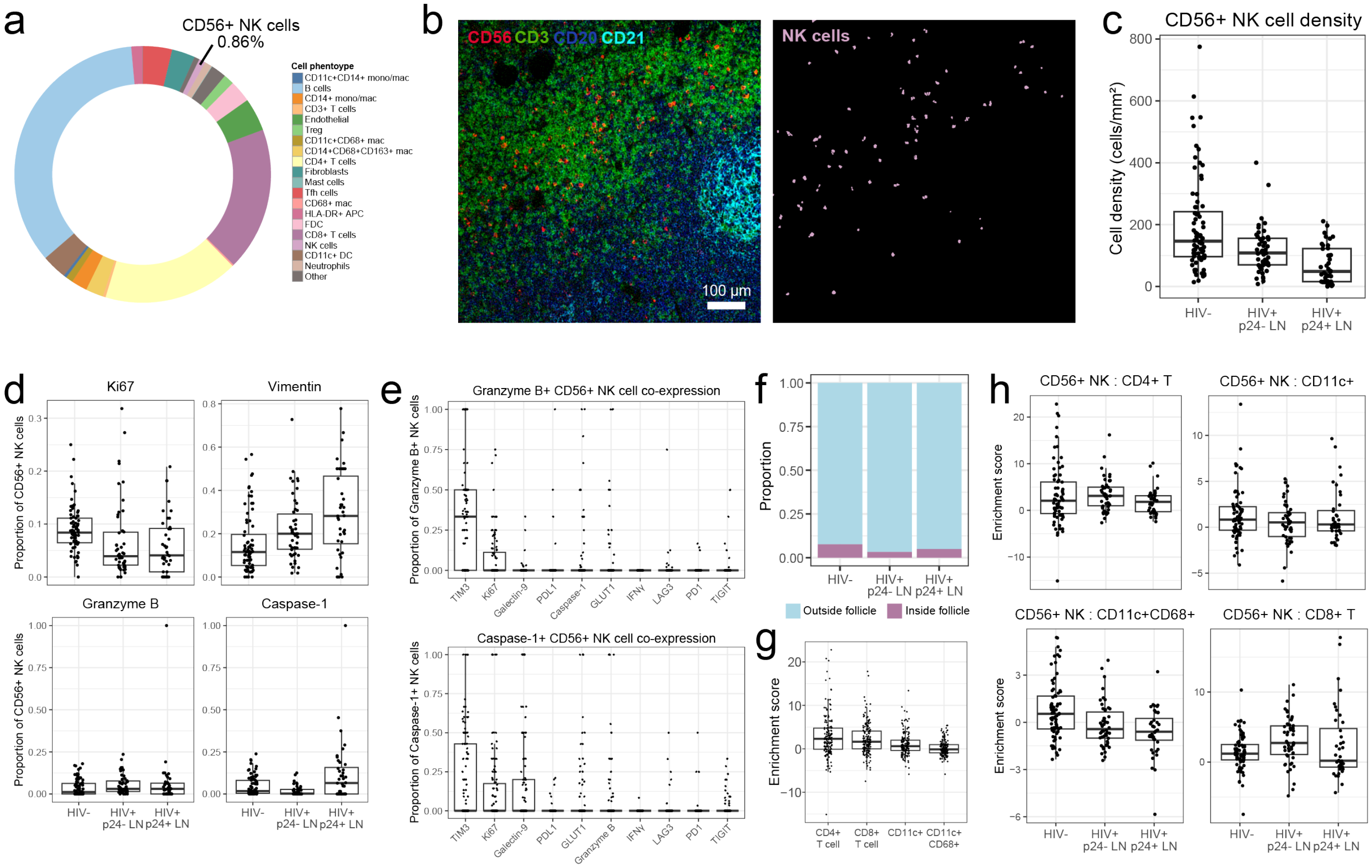


**Supplementary Figure 7: Features of CD56+ NK cells in LNs**

(a) Proportion of cell phenotypes identified in the cohort. (b) Representative MIBI-TOF image (left) and CD56+ NK cell detection (right). (c) Comparison of CD56+ NK cell density between the donor groups. (d) Comparison of functional marker expression in CD56+ NK cells. (e) Proportion of granzyme B+ (top) and caspase-1+ (bottom) CD56+ NK cells that express other functional markers. (f) Proportion of CD56+ NK cells that are inside or outside the follicles. (g) Enrichment scores of CD56+ NK cells with other cell populations. (h) Comparison of CD56+ NK cell enrichment scores across donor groups.

Supplementary Figure 8

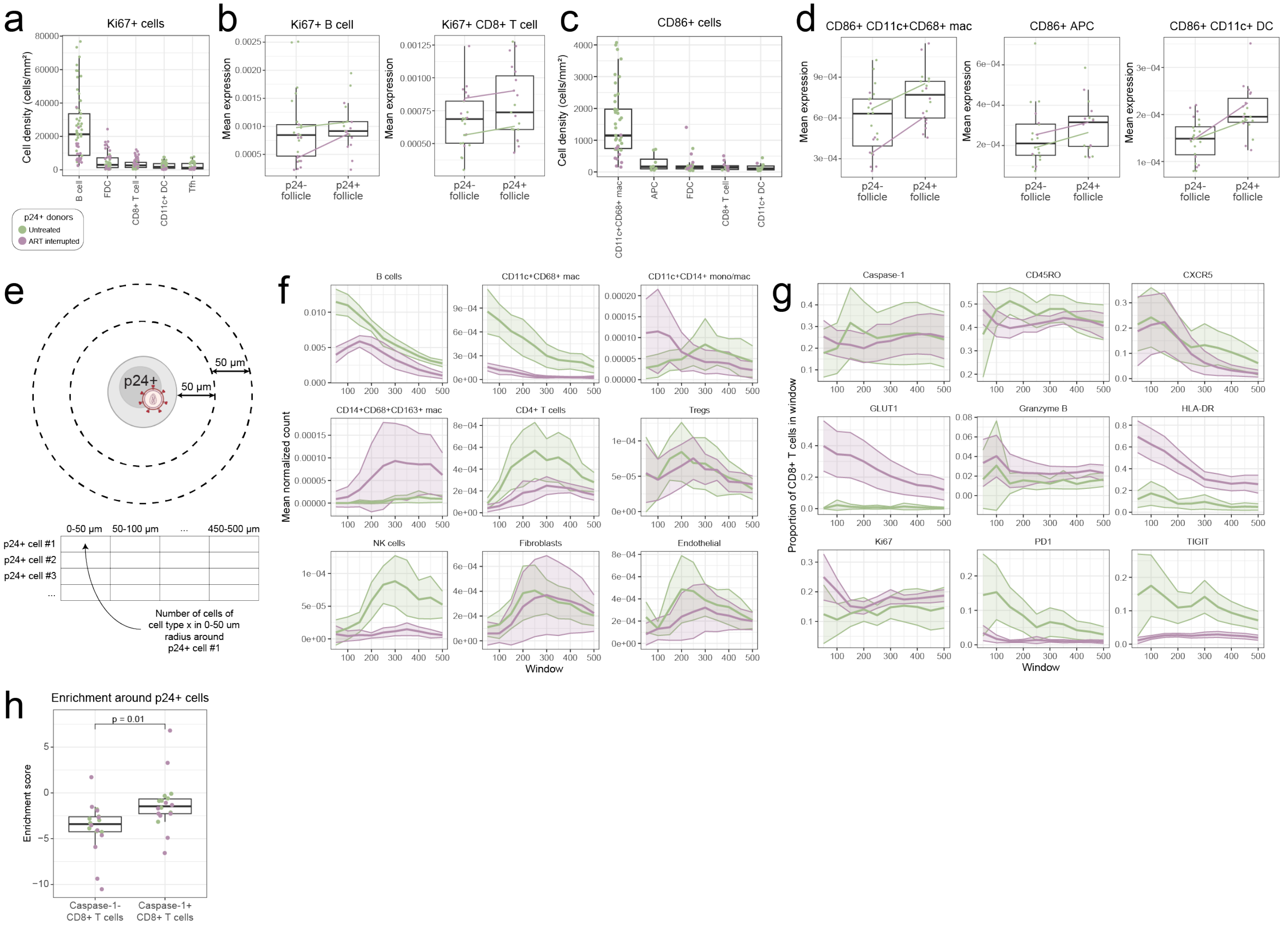


**Supplementary Figure 8: Tissue microenvironments in the neighborhood of active viral replication in untreated versus treatment interrupted PWH**

(a) Breakdown of Ki67+ cells by cell density. Colors correspond to different donors based on treatment status. Each dot is one follicle. (b) Ki67 expression in B cells in CD8+ T cells. Each dot is one follicle. The lines connect the medians of the two groups. (c) Breakdown of CD86+ cells by cell density. (d) CD86 expression in myeloid cell populations. (e) Overview of p24+ cell distance analysis. For each p24+ cell, the proportion of each cell type was enumerated in increasing distance windows of 50 μm from the center cell. (f) Density of each cell type or (g) functional marker positive CD8+ T cells in increasing distance radii around p24+ cells. Lines represent the mean and shaded areas represent 95% confidence intervals. (h) Comparison of enrichment score between caspase-1- and caspase-1+ CD8+T cells with p24+ cells. Each dot is one FOV.

#### Supplementary Figure 9

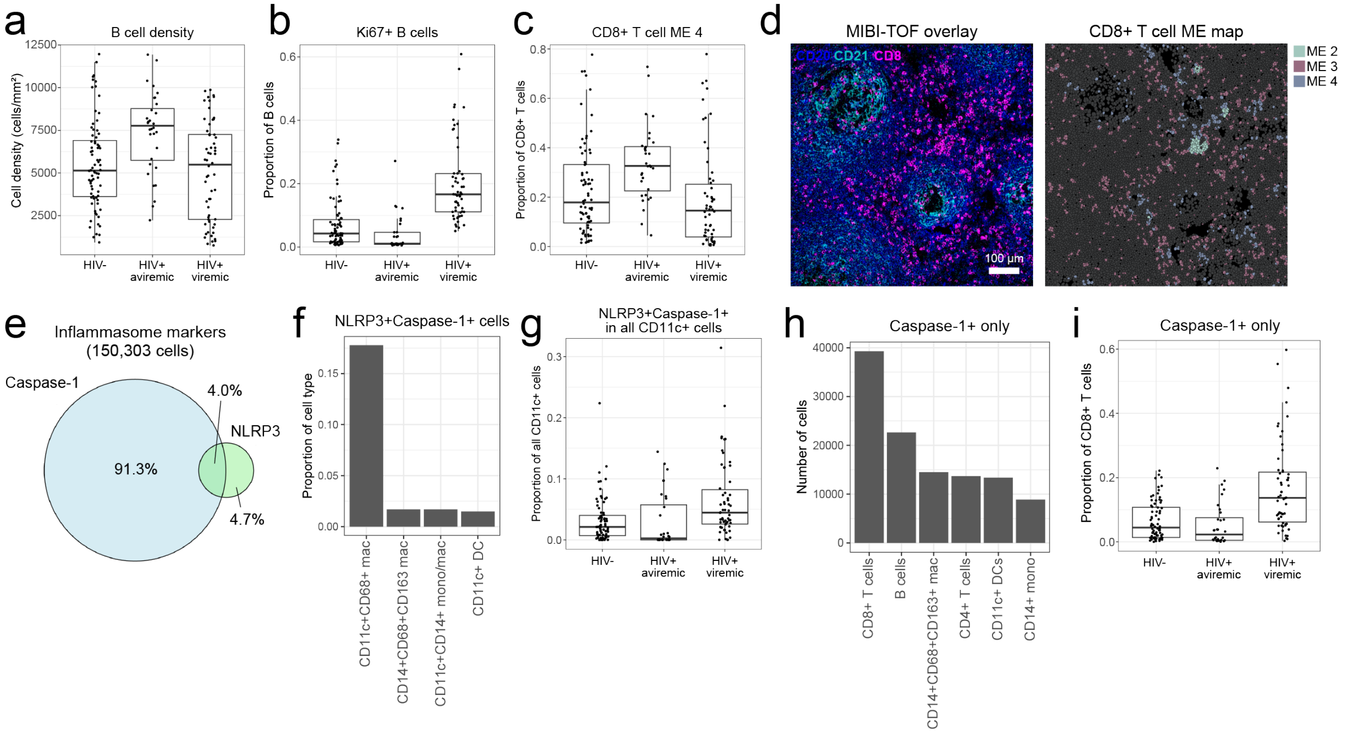


**Supplementary Figure 9: Features of LNs from aviremic donors and inflammasome marker distribution**

(a) Comparison of B cell density between HIV-, aviremic, and viremic groups, defined by virus detectable in the plasma. (b) Proportion of B cells expressing Ki67. (c) Comparison of CD8+ T cells belonging to ME cluster 4 across viremia groups. (d) Representative MIBI-TOF image (left) and corresponding ME map (right) where the colors correspond to cluster number. Gray cells are cells that are not CD8+ T cells. (e) Venn diagram of all cells expressing inflammasome markers caspase-1 or NLRP3. (f) Breakdown of NLRP3+caspase-1+ cells. (g) Proportion of all CD11c+ cells expressing NLRP3 and caspase-1. (h) Breakdown of cells that are caspase-1+NLRP3-. (i) Comparison of proportion of CD8+ T cells that express caspase-1.
